# Cross-scale visualization of RNA dynamics in whole plants using an RNA-triggered reporting system

**DOI:** 10.1101/2025.03.03.641157

**Authors:** Jiuyuan Bai, Yazhou Chen, Jiayu Zhang, Hongpei Ren, Shan Pan, Yiran Tao, Miaomiao Lei, Mengyue Dong, Sha Liu, Jia Yi, Fang Chen, Jiang Li, Chunhai Fan, Yun Zhao

## Abstract

Real-time visualization of RNA dynamics across spatial scales in living plants is desirable yet remains challenging, hindered by the absence of imaging tools that simultaneously offer high sensitivity and deployability in whole living plants. Here, we engineered a modular, RNA-triggered fluorescence (RTF) reporter system *de novo* for spatiotemporal imaging of RNA from cellular to whole-plant scales *in vivo*. This system integrates three functional modules: a target-specific allosteric RNA switch, a degradable adapter for background suppression, and a fluorescent reporter. All components are stably expressed from a single vector and assemble *in vivo*, enabling uniform deployment throughout entire plants. The modified RTF (RTFst) achieves single-molecule sensitivity (signal-to-noise ratio >60) for dynamic tracking of developmentally regulated, tissue-specific, circadian, and stress-responsive mRNAs. Crucially, it enables the first direct, real-time observation of graft-transmissible mRNA trafficking across tissue scales and cross-kingdom transfer of the aphid-secreted long non-coding RNA *Ya1* to plant cells. RTF establishes a versatile platform for studying gene regulation, signaling, and transcript mobility in plants, with broad applicability in synthetic biology and crop improvement.

## Introduction

RNA accumulation is highly dynamic and deeply intertwined with the physiological state of the whole organism ^1–3^, however, capturing these dynamics at the scale of whole living systems remains a significant challenge in biology.

Whole living organisms are important for understanding RNA expression dynamics ^4^, which reflect the complex regulatory networks governing growth, development, and adaptation to environmental changes. Within individual cells, RNA expression is shaped not only by the local cellular conditions but also by systemic signals originating from distant tissues ^5–9^. These processes are further modulated by external environmental signals that directly act on the organism as a whole. Moreover, RNAs are often trafficked between tissues, underscoring their dynamic nature and functional importance ^5^. Intriguingly, living organisms frequently acquire RNAs from other organisms that belong to entirely different biological kingdoms, with transfer of insect RNAs known to occur into plants on which the insect feeds ^10–13^. Those RNAs function within the recipients, mediating intimate interspecies relationships ^14^. Profiling such RNA dynamics across time and space at the scale of whole living organisms is crucial for understanding gene regulation and ecological interactions within interconnected physiological and environmental contexts. However, achieving this is difficult, in particular for biotic interactions.

Over the past few decades, numerous single-molecule RNA tracking techniques have been developed to visualize RNA dynamics in living systems, largely in animal systems. These approaches generally fall into two major categories. The first relies on genetically encoded tags appended to target RNAs, such as fluorescent RNA aptamers^15, 16^ or RNA motifs such as MS2 that recruit exogenous reporters (e.g., fluorescent proteins) ^1^. While these systems have enabled high-resolution imaging, the introduction of RNA tags can perturb the native properties of the RNA, including its abundance, expression dynamics, stability, subcellular localization, and mobility ^17–20^. A second category comprises methods that detect endogenous RNAs via trans-binding probes, either chemically synthesized or genetically encoded. This group includes molecular beacons^21^, split probes^22^ and CRISPR-based reporters^23–26^. Although these methods, in principle, detect unmodified RNAs, they are often limited by high background fluorescence, insufficient spatial resolution for single-molecule level, and inefficient or non-uniform probe distribution across large, complex live tissues^27^. Recently, a split ribozyme system was adapted for *in vivo* RNA imaging in plants ^28^. While promising, its reliance on ribozyme-mediated GFP transcript reassembly followed by translation introduces a temporal lag incompatible with real-time RNA tracking. Moreover, ribozyme splicing efficiency is highly variable in plant cells due to the complex intracellular environment, further limiting its applicability for visualizing long-distance RNA trafficking at single-molecule resolution *in vivo*.

To address these limitations, we developed an RNA switch-controlled RNA-triggered fluorescence system (RNA switch–RTF), enabling high-resolution and sensitive visualization, monitoring, and quantification of RNA dynamics in living plants. Using RNA switch–RTF, we tracked *in vivo* expression dynamics of multiple conditionally expressed RNAs in the model plant *Arabidopsis thaliana* at subcellular and single-cell resolution throughout the entire plant. RNA switch–RTF demonstrated cell-to-cell trafficking of *TRANSLATIONALLY CONTROLLED TUMOR PROTEIN 1* (*TCTP1*) mRNAs ^29^ in *A. thaliana* and systematic migration of aphid-derived long non-coding RNA *Ya1* ^30^ in its leaves. This approach provides a robust tool for studying RNA localization and movement *in vivo*, offering critical insights into RNA function in plant development and physiology, and ecological interactions.

## Results

### *De novo* design of an RNA-triggered fluorescence (RTF) reporter system

We sought to develop a fluorescent reporter system to visualize specific RNA molecules in whole plants. In this system, GFP fluorescence would be detectable only when the target RNA is present. This requires GFP to be constitutively degraded in plants in the absence of the target RNA, with degradation inhibited upon the presence of the target (Fig. 1A). Theoretically, constitutive GFP degradation can be achieved by tagging it with degron sequences^31^— minimal motifs that recruit 26S proteasomes to target proteins.

**Fig. 1.**
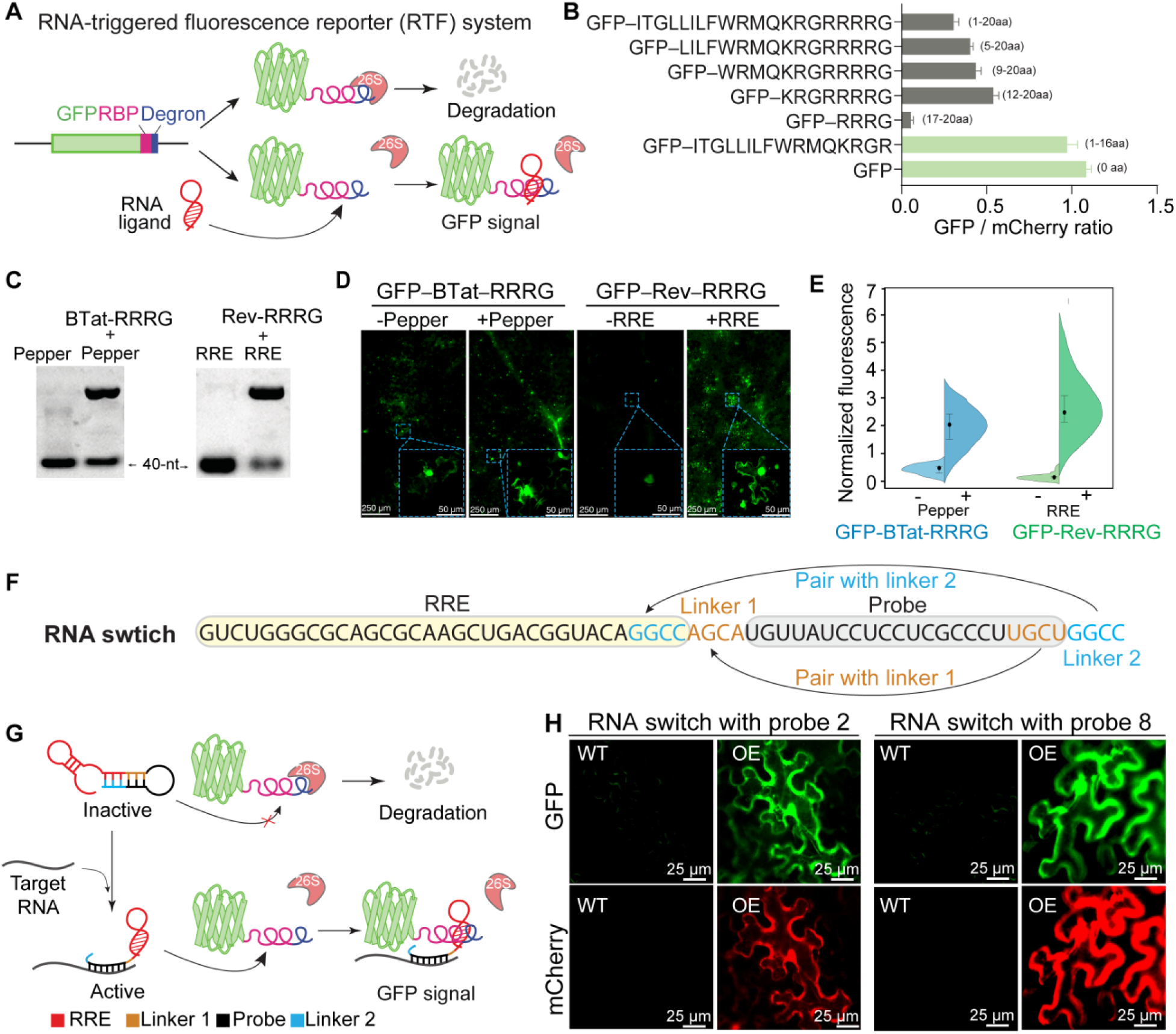
Design of the RNA switch-controlled RTF system. (**A**) Schematic of the RTF reporter system. The fusion protein GFP–RBP–Degron undergoes degradation in the absence of RNA ligands. Binding of the RNA ligand to the RBP in the fusion protein blocks accessibility to the 26S proteasome, resulting in GFP signal accumulation. (**B**) Assessment of the degradation ability of putative degron sequences. Degradation efficiency was evaluated based on the GFP/mCherry fluorescence intensity ratio. mCherry and GFP tagged with the putative degron sequence were co-expressed using a 2A self-cleaving peptide system (Fig. S1A). The mCherry–2A–GFP–degron construct was cleaved at the 2A site, producing separate mCherry and GFP–degron proteins. (**C**) Binding abilities of BTat–RRRG and Rev–RRRG to their specific RNA ligands, Pepper and RRE, respectively. (**D** and **E**) GFP fluorescence signals in the absence and presence of specific RNA ligands when tagged with BTat–RRRG or Rev–RRRG. (**F**) Design of the RNA switch. Sequences shaded in yellow comprise the RRE and in gray are the probe. The sequences in brown are linker 1 and those in blue are linker 2. (**G**) Diagram of the RNA switch-controlled RTF reporter system. In the absence of the target RNA, the RNA switch adopts an inactive state through base pairing between linkers and their complementary nucleotides in the probe and RRE. When the target RNA is present, the probe hybridizes with the target, unfolding the RNA switch and enabling it to bind the RBP in GFP–RBP–RRRG. (**H**) RNA switches containing probes 2 and 8 that target *mCherry* RNA, are inactive in *N. benthamiana* WT plants but active in transgenic lines expressing *mCherry*.

Although several degrons have been characterized in plants ^32, 33^, their relatively large size limits their suitability for adaptation into compact RNA-binding peptides. Therefore, we first identified degrons in leaves of the model plant *Nicotiana benthamiana* that is amenable to and convenient for transient and stable transformation. By comparative proteomics, 17 proteins strongly accumulated only in leaves treated with the proteasome inhibitor MG132, compared to mock-treated controls (Table S1, Fig. S1A–C). We fused 20 amino acids at the C-terminus of these 17 proteins to the C-terminus of GFP and tested their ability to promote GFP degradation relative to an mCherry internal control (Fig. S1D, E). Twenty residues at the C terminus of Niben261Chr01g0008021.1 (encoding a predicted subtilisin-like protease SBT6) enhanced GFP degradation (Fig. 1B, Fig. S1E). Further analysis identified the final four residues of this protein, RRRG, as a degron capable of efficiently promoting GFP degradation (Fig. 1B, Fig. S1F).

To enable GFP degradation precisely in response to the target RNA, we chose six well-characterized RNA-binding peptides (RBP) ^34–39^ (Table S2), each with specificity for RNA cognate aptamer. Fusion of the RRRG degron to the C terminal of these RBPs did not affect their capacity to bind RNA (Fig. 1C, Fig. S1G). To assess the functionality of these RBPs fused with RRRG, we next tested the abundance of GFP tagged with RBP–RRRG in the presence and absence of specific RNA aptamers. Of these, GFP–BTat–RRRG and GFP–Rev–RRRG accumulated in *N. benthamiana* leaves when exposed to RNA aptamers Pepper and RRE, respectively (Fig. 1D, Fig. S1H). GFP–Rev–RRRG showed superior performance, with strong GFP fluorescence in the presence of RRE, while no signal was detectable in its absence by fluorescence imaging (Fig. 1E) and western blotting (Fig. S1, H and I).

In sum, we successfully developed RNA-triggered fluorescence (RTF) by manipulating GFP degradation to visualize the presence of an RNA ligand in plants.

### Engineering of RNA switches to control GFP degradation for use with RTF

The RNA aptamer RRE binds to Rev and inhibits the recruitment of the proteasomes to the degron on the GFP–Rev–RRRG chimeric protein (Fig. 1D, E). Building on this, we engineered RNA switches capable of controlling GFP degradation in the presence of specific target RNA molecules. To achieve this, we constructed RNA switches based on RRE (39 nt) that incorporate three functional elements: a probe sequence complementary to the target RNA and two linker sequences that hybridize with the RRE and the probe, respectively (Fig. 1F).

Theoretically, in the absence of the target RNA, the RNA switch adopts an inactive conformation whereby the linkers hybridize with the RRE and the probe. This folding prevents the RRE from binding to the degron, leading to GFP degradation (Fig. 1G). Conversely, when the target RNA is present, the RNA switch transitions to an active conformation through base pairing between the probe and the target. This conformational change allows RRE to bind to the degron, leading to GFP accumulation (Fig. 1G).

To evaluate the functionality of the RNA switch, we designed various combinations of probes and linker sequences, and tested their ability to regulate GFP levels in the presence and absence of the target RNA. Using a stable-transgenic *N. benthamiana* line expressing *mCherry* under the 35S promoter, we designed 12 distinct probes to target different regions of *mCherry* mRNA (Fig. S2A, Table S3). The RNA switch plasmid and the GFP–Rev–RRRG plasmid in *Agrobacterium tumefaciens* were co-infiltrated into *N. benthamiana* leaves of both *mCherry* and wild-type (WT) backgrounds. Probes 2 and 8 demonstrated functionality (Fig. S2, B and C). In WT plants, these probes maintained the RNA switch in an inactive state, as evidenced by the absence of detectable GFP fluorescence (Fig. 1H). In *mCherry* plants, however, these probes enabled the RNA switch to bind the degron, resulting in strong GFP fluorescence (Fig. 1H).

RNA switches with probes 2 and 8 adopted a dumbbell-shaped structure (Fig. S2A), which appears critical for functionality. The switch incorporated with probe 2 has a shorter stem sequence and a larger bottom loop, enhancing its efficiency. Other structural configurations failed to regulate GFP signals effectively (Fig. S2, A to C), underscoring the importance of the dumbbell-shaped conformation for RNA switch performance.

These engineered RNA switches effectively control the RTF system and enable precise regulation of GFP degradation in response to specific RNA molecules.

### Optimization of the RNA switch-controlled RTF system in plants

To evaluate the RNA switch–RTF system in plants, we constructed a plasmid containing an RNA switch targeting the Arabidopsis *CHORISMATE MUTASE 3* (*CM3*) mRNA driven by the *U6* promoter, and GFP–Rev–RRRG under the control of the *UBQ10* promoter (Fig. 2A). These expression modules were introduced into both Arabidopsis wild-type (WT) and *cm3* mutant plants (Fig. 2A). In WT plants, strong GFP signals were observed, but *cm3* mutant plants also produced weak but detectable GFP signals (Fig. 2B), indicating that the system displayed some degree of signal leakage. The leakage was also confirmed by the observation of GFP in WT plants expressing RNA switch–RTF that were treated with the RNA synthesis inhibitor Actinomycin D (Fig. S3, A and B).

**Fig. 2.**
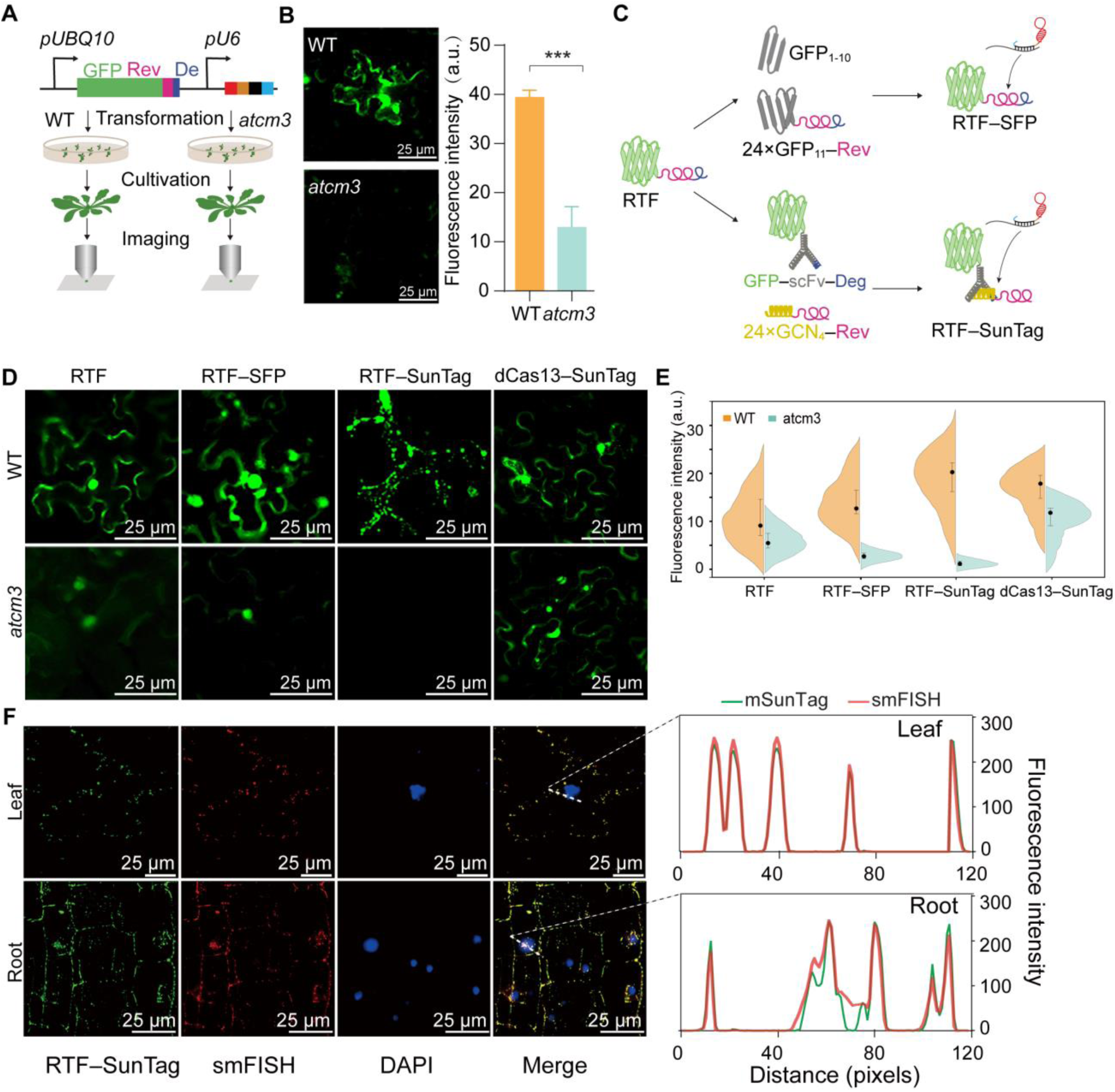
Optimizing the RNA switch-RTF system in plants. (**A**) Schematic representation of introducing the RNA switch–RTF system into Arabidopsis. The RTF system, driven by the *UBQ10* promoter, and the RNA switch targeting *CM3* mRNA, driven by the *U6* promoter, are on the same plasmid. The plasmids were introduced into *A. thaliana* WT and *cm3* mutant plants. (**B**) GFP fluorescence signals in WT and *cm3* mutant plants. Left: confocal images. Right: fluorescence intensity. (**C**) Optimizing the GFP reporter by replacing it with a split fluorescent-protein (SFP) approach and SunTag. (**D** and **E**) Fluorescence signals in *A. thaliana* WT and *cm3* mutant plants genetically introduced with RNA switch-controlled RTF, RTF–SFP, RTF–SunTag, and dCas13–SunTag. Confocal images are shown in D and quantification of fluorescence intensity is shown in E. (**F**) Localization of *CM3* mRNA visualized using RNA switch–RTF–SunTag and smFISH. Cy3-labeled probes specific to *CM3* mRNA were used for smFISH. The fluorescence intensity of labeled regions in the merged images is shown on the right. Green lines represent GFP fluorescence intensity, while red lines indicate Cy3 fluorescence intensity.

To address this issue, we incorporated two additional strategies: a split fluorescent protein (SFP) approach ^3^ and SunTag ^40^ (Fig. 2C). In the SFP strategy, GFP–Rev–RRRG was replaced with two modules: GFP_1–10_ and 24×GFP_11_–Rev. With SunTag, GFP–Rev–RRRG was also replaced with two components: full-length *GFP* fused to a single-chain antibody (scFv) tagged with the RRRG degron and 24×GCN4 tagged with Rev at its N-terminus. These systems, combined with RNA switches targeting *CM3* mRNA, were introduced into WT and *cm3* mutant plants. RTF–SunTag plants expressed an extremely low GFP signal in *cm3* mutant plants compared to the RTF, RTF–SFP systems (Fig. 2, D and E, Fig. S3C), likely due to degradation of unbound GFP.

We also compared the RTF–SunTag with dCas13–SunTag system, a robust RNA imaging tool in mammalian cells. However, in *cm3* mutant, dCas13–SunTag system produced high background fluorescence, resulting in false positives and difficult to interpret the signal in WT plants (Fig. 2, D and E). This background is likely due to large vacuoles in plant cells, which compress the cytosol in a thin peripheral layer, concentrating even minor cytosolic fluorescence and thereby amplifying background noise^41^. In contrast, the RTF–SunTag system mitigated this issue by incorporating a degron we screened (Fig. 2C), which significantly reduced background and yielded a 12-fold increase in signal-to-background (S/B) ratio compared to the dCas13– SunTag system (Fig. 2, D and E).

To assess the accuracy of the RNA switch–RTF–SunTag system, we compared the localization of *CM3* mRNA indicated by RTF–SunTag with that determined by smFISH using a series of Cy3-labeled probes targeting *CM3* mRNA. Colocalization of GFP and Cy3 signals was observed in the leaves and roots of transgenic Arabidopsis plants containing RNA switch–RTF– SunTag (Fig. 2, F to G, Fig. S3, E and F), confirming its accuracy in detecting *CM3* mRNA localization.

In summary, the RNA switch–RTF system, optimized with SunTag and hereafter referred to as RNA switch–RTFst, accurately localizes the target RNA within plant cells, providing a robust and reliable tool for studying RNA dynamics and their subcellular localization.

### RNA subcellular localization is unaffected by RNA switch–RTFst

RNA function is often closely tied to subcellular localization ^42^. We designed three experiments to assess whether RNA switch–RTFst influences RNA localization. First, we compared *CM3* subcellular localization resulting from the RNA switch–RTFst system with those from the mCherry–MCP–MS2 system. In the latter system, repeated *MS2* sequences were fused to the 3’ UTR of *CM3* mRNA, allowing the MS loops on the hybrid RNA to capture MCP from the mCherry–MCP protein (Fig. 3A). *CM3* mRNAs localized in both nuclei and the cytoplasm (Fig. 3A). With RNA switch–RTFst, probe sequences targeting the 5’ UTR, coding sequence, and 3’ UTR of *CM3* were designed (Table S5). Probes targeting the coding sequence showed identical subcellular localization patterns to those observed with the mCherry–MCP–MS2 system (Fig. 3A, Fig. S3, G and H). Probes targeting the 5’ UTR should be used with caution because these tend to detect fewer RNA molecules per cell.

**Fig. 3.**
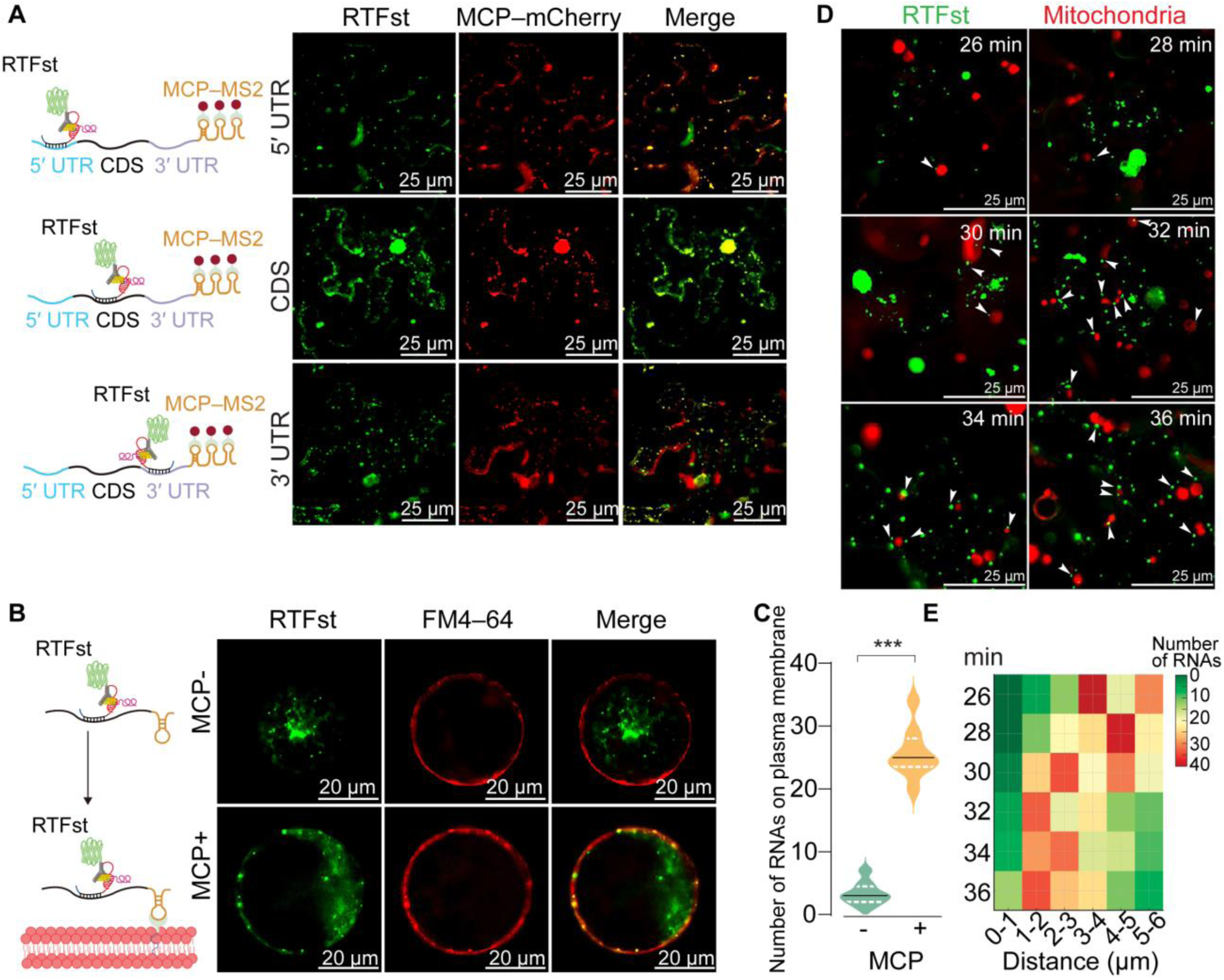
RNA localization is unaffected by the RNA switch–RTFst complex. (**A**) RNA localization determined by RNA switch–RTFst is comparable to that assessed by MCP– MS2. The *MCP–MS2* plasmid contains 24× *MS2* loops fused to sequences coding for the 5′ UTR, CDS, and 3′ UTR of *CM3* mRNA, driven by the *UBQ10* promoter, as well as *MCP* sequences driven by the CaMV *35S* promoter. The MCP–MS2 plasmid and the RNA switch–RTFst plasmid with probes targeting the 5′ UTR, CDS, or 3′ UTR were transiently co-expressed in *N. benthamiana* WT leaves. (**B**) The RNA switch–RTFst complex targeting *CM3* mRNA did not obviously affect RNA localization. For the MCP– condition, the RNA switch–RTFst plasmid with a probe targeting the *CM3* coding sequence and a plasmid expressing *CM3*–*MS2* mRNA were co-expressed in *N. benthamiana* mesophyll protoplasts. For the MCP+ condition, the RNA switch– RTFst plasmid, and the *CM3*–*MS2* mRNA plasmid containing the *MCP* sequence tagged with CAAX (a membrane targeting motif) were co-expressed in *N. benthamiana* mesophyll protoplasts. Cell membranes were stained with FM4-64. (**C**) Quantification of RNAs localized to the cell membrane. Statistical significance (\*\*\**p* < 0.001) was calculated using two-tailed unpaired Student’s *t*-test. (**D**) *VDAC3* RNA with the RNA switch–RTFst complex localizes to mitochondria. Green dots represent *VDAC3* RNAs visualized by RNA switch–RTFst, while red dots indicate mitochondria stained with Red CMXRos. White arrows highlight *VDAC3* RNAs in proximity to mitochondria. (**E**) Time-dependent proximity of *VDAC3* RNAs to mitochondria

Second, the RNA switch-controlled RTFst system with a probe targeting *CM3*, and *CM3*– *6×MS2* RNA was co-expressed in protoplast cells obtained from *N. benthamiana* leaves. GFP signals revealed that *CM3*–*6×MS2* RNA was predominantly localized in the cytoplasm and was nearly undetectable at the plasma membrane (Fig. 3, B and C). However, when MCP tagged with CAAX (a membrane-targeting motif) ^43^ was introduced, GFP signal along the membrane increased (Fig. 3, B and C). This change was attributed to the binding of MS loops on *CM3*–*6×MS2* RNA to MCPs at the membrane. These results suggest that the localization of *CM3*–*6×MS2* RNA, even when complexed with the RNA switch–RTFst, remained unaffected.

Third, we evaluated the RNA switch–RTFst system using the mitochondria-localized RNA *VDAC3* (*VOLTAGE-DEPENDENT ANION CHANNEL 3*), which has been reported to reside in the mitochondria of *N. benthamiana* cells ^44^. RNA switch–RTFst and *VDAC3* RNA were co-expressed in *N. benthamiana* leaves. Over time, the GFP signal became increasingly proximal to mitochondria (Fig. 3, D and E). This progressive colocalization suggested that *VDAC3* RNA was unaffected by the system and remained associated with its expected subcellular compartment, the mitochondria.

RNA switch–RTFst does not detectably alter the subcellular localization of RNA in plants, maintaining native localization patterns of at least these target transcripts.

### RNA switch–RTFst is suitable for real-time monitoring of RNA dynamics in living plants

To enable real-time monitoring of RNA dynamics in living plants, we applied the RNA switch–RTFst to a set of transcripts representing diverse regulatory contexts. Specifically, we selected *TUB5* (*TUBULIN β-5*) for development-dependent expression ^45^, *SMB* (*SOMBRERO*) for tissue-specific expression ^46^, *CCA1* (*CIRCADIAN-CLOCK-ASSOCIATED 1*) for circadian regulation ^47^, and *CIPK11* (*CBL-INTERACTING PROTEIN KINASE 11*) for stress responsiveness, particularly to cadmium (Cd) ^48^ (Table S6). RNA switches targeting the coding sequences of those RNAs were introduced, along with RTFst, into *A. thaliana* wild-type plants.

*TUB5* mRNA is ubiquitously expressed in *A. thaliana*, with lower levels in vegetative tissues and higher levels in stems and reproductive tissues. GFP signals were detected across all examined tissues (Fig. 4, A and B). During development, signals remained relatively stable in roots but became stronger in flowers (Fig. 4, A and B, Fig. S4, A and B). RNA switch–RTFst showed no obvious effect on the expression, turnover, or translation of *TUB5* mRNA, or on overall ubiquitination patterns (Fig. S5 A to E).

**Fig. 4.**
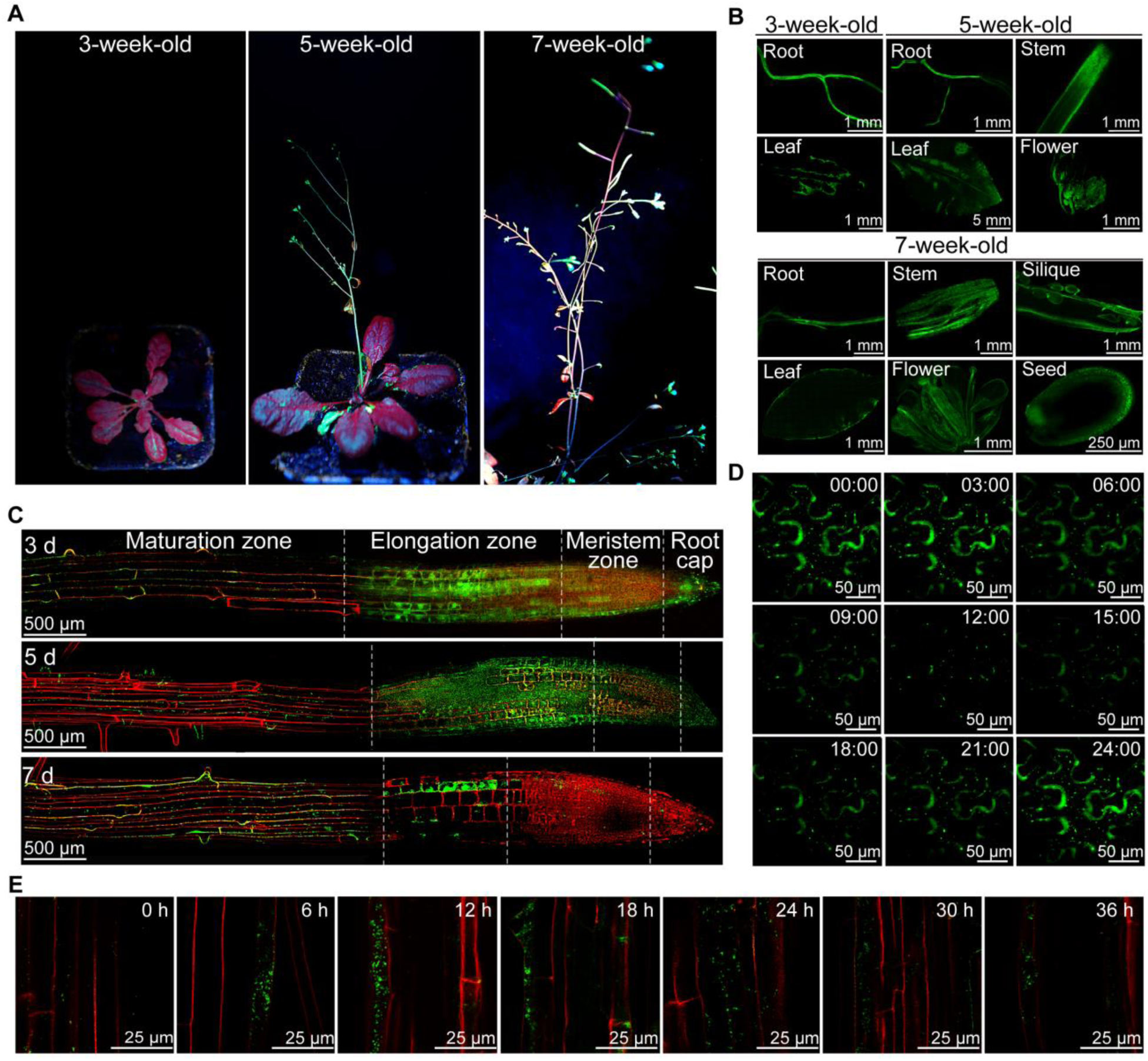
RNA switch–RTFst enables monitoring dynamic changes in RNA expression at different spatial scales. (**A** and **B**). Developmental stage- (A) and tissue-dependent (B) expression levels of *TUB5*. Transgenic *A. thaliana* plants expressing RNA switch–RTFst targeting *TUB5* RNAs were analyzed. (**C**) Tissue-specific expression patterns of *SMB*. Root tips from transgenic *A. thaliana* plants expressing RNA switch–RTFst targeting *SMB* RNAs were harvested. Cell walls (red) were stained with propidium iodide. (**D**) Circadian-regulated expression patterns of *CCA1*. Leaves from 7-week-old transgenic *A. thaliana* plants expressing RNA switch–RTFst targeting *CCA1* RNAs were collected every 3 h, and GFP signals were observed. (**E**) Cd-induced expression patterns of *CIPK11*. Seven-week-old transgenic *A. thaliana* plants expressing RNA switch–RTFst targeting *CIPK11* RNAs were treated with 50 μM Cd. Roots were harvested every 6 h post-treatment and GFP signals were recorded. Cell walls (red) were stained with propidium iodide. Green fluorescence represents target RNAs visualized by RNA switch–RTFst.

*SMB* was exclusively expressed in the specialized zones of roots, mainly in the elongation zone of roots in 3- and 5-d-old seedlings, with a gradual increase in expression observed in the maturation zone of 7-d-old seedlings (Fig. 4C, Fig. S4, C and D). *CCA1* is a circadian-regulated RNA that exhibits time-of-day-specific expression ^47^. Consistent with this, here its abundance displayed dynamic changes during the morning and evening phases (Fig. 4D, Fig. S4, E to F).

*CIPK11*, a Cd-responsive mRNA ^48^, was up-regulated in roots 12 h after Cd treatment, followed by gradual down-regulation at 18 h (Fig. 4D, Fig. S4, H to I). These dynamic expression patterns identified by RNA switch–RTFst were validated by qRT–PCR (Fig. 4, E and F, Fig. S4, G and J).

Altogether, RNA switch–RTFst is a powerful tool to visualize and monitor the *in vivo* dynamics of RNA expression in living plants, offering insights into development, tissue specificity, circadian regulation, and stress responses.

### RNA switch–RTFst monitors the movement of mobile RNAs in living plants

In plants, transcripts can move to distant tissues, potentially acting as systemic signals regulating development and growth ^49^. *TCTP1* mRNA has long-distance mobility, traveling from the shoot to the root through the phloem ^29^. To track this movement, we expressed the *TCTP1*-targeted RNA switch–RTFst system in *tctp1* mutants (*tctp1*–*RTFst*) (Fig. 5A). Grafting experiments were then conducted using wild-type shoots as the scion and *tctp1*–*RTFst* roots as stock (Fig. 5A). At 72 h post-grafting, GFP signals progressively intensified, particularly at 5 mm below the graft junction (Fig. 5, B and C), and stronger GFP signals accumulated in root sections further from the junction over time (Fig. 5, B and C). Time-lapse imaging confirmed the movement of *TCTP1* mRNA across the graft junction (Fig. 5, B and D, Movie 1). Upon reaching the root, *TCTP1* mRNA is known to influence lateral-root formation and regulate root architecture (*32*). Root length in grafted plants was comparable to that of WT and statistically significantly longer than in *tctp1* mutants (Fig. 5D), suggesting that WT-derived *TCTP1* mRNA compensated for the loss of function in the roots. This compensation was unaffected by RNA switch–RTFst expression in the roots.

**Fig. 5.**
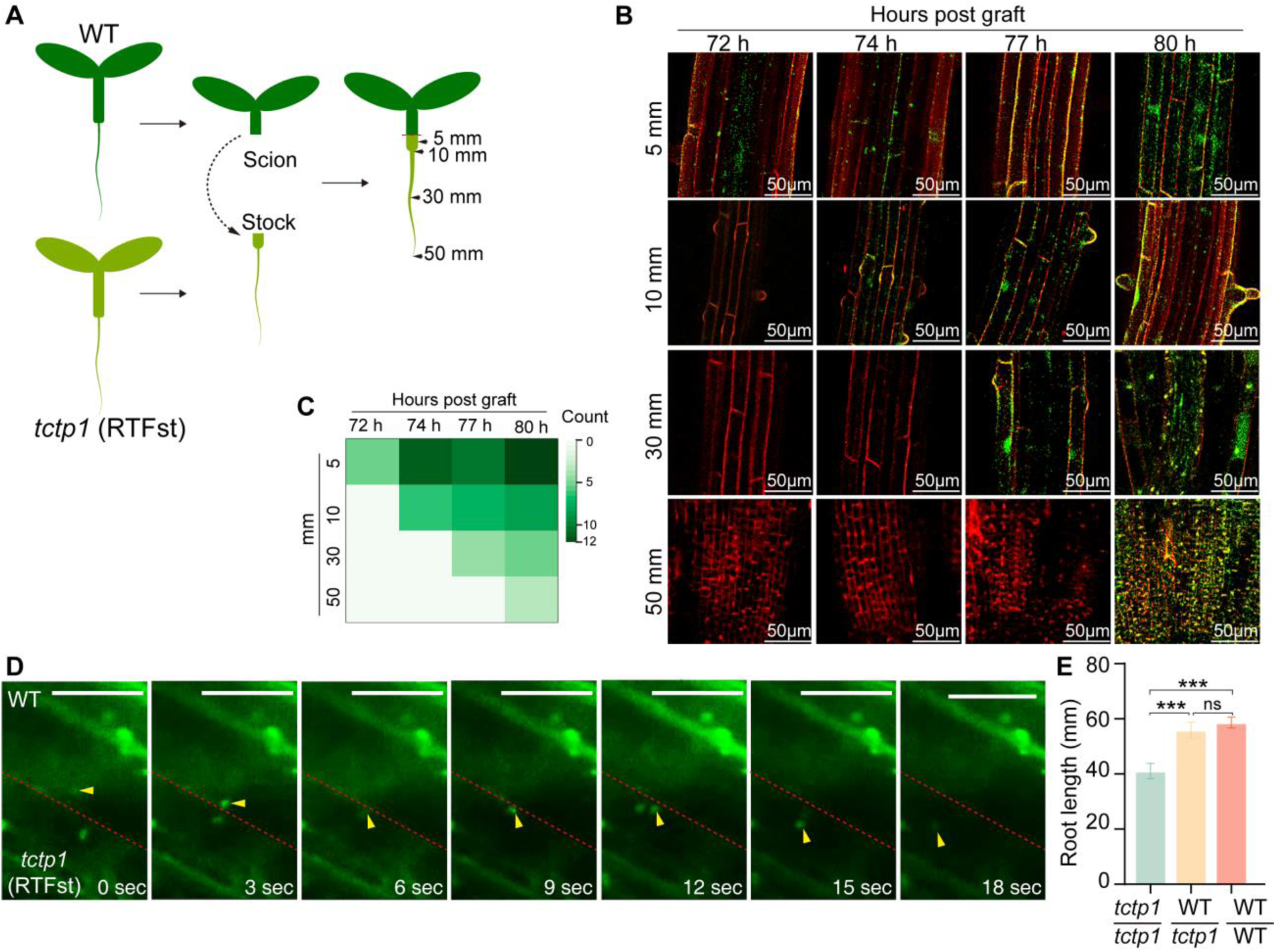
RNA switch–RTFst visualized the real-time cell-to-cell trafficking of *TCTP1* mRNA. (**A**) Schematic of hypocotyl grafting. One-week-old Arabidopsis seedlings of similar sizes and with evenly elongated hypocotyls were cut in the upper half of the hypocotyl with a sterilized razor blade. WT shoots were used as scions and *tctp1*–*RTFst* roots as stocks. The red dashed line indicates the graft junctions. The black arrowhead indicates the positions of 5 mm, 10 mm, 15 mm, and 20 mm below the graft junctions. Grafted plants were grown vertically under short-day conditions. (**B**) Representative imaging of *TCTP1* RNA in different seedlings positions after grafting. Cell walls (red) were stained with propidium iodide. (**C**) Quantification of grafted plants with fluorescence signals at various times post-grafting. Twelve grafted seedlings were imaged for each selected position below the graft junction. (**D**) Real-time fluorescence imaging of cell-to-cell movement of *TCTP1* RNA in root cells of grafted plants. See the **video S1** for details. The red dashed line indicates the cell boundary and the yellow arrowhead indicates *TCTP1* RNA. Scale bar = 25 μm. (**E**) Root length of *tctp1* mutants, grafted plants, WT plants (n = 6, mean ± S.E., *** *P* < 0.001, one-way ANOVA with multiple comparisons and a post-hoc test).

### RNA switch–RTFst can visualize the aphid-secreted lncRNA *Ya1* in plants

Beyond endogenous RNA movement within plants, RNAs can also be transferred between organisms from distinct biological backgrounds ^50^, with the green peach aphid *Myzus persicae* translocating the lncRNA *Ya1* into host plants via feeding ^30^. To investigate the mobility of this aphid-secreted RNA, we designed a *Ya1*-targeted RNA switch–RTFst and transiently expressed it in *N. benthamiana* leaves. When co-expressed with *Ya1*, GFP signals were detected in the cytoplasm and nuclei of infiltrated cells (Fig. 6A). In leaves where the construct was expressed and subsequently fed on by aphids, GFP signals accumulated at the feeding sites (Fig. 6A). Furthermore, when the system was expressed in two separate infiltration sites on the same leaf, with one co-expressing *Ya1*, GFP signals were observed at the distal site, even in the absence of direct *Ya1* expression (Fig. 6B, Movie 2). To visualize the dynamics of *Ya1* translocation, *N. benthamiana* leaves expressing *Ya1*-targeted RNA switch–RTFst were exposed to aphid feeding and monitored by time-lapse imaging. After 6 h of continuous feeding, GFP signal was detectable and intensified over time, while control leaves expressing RNA switch–RTFst but without aphid feeding exhibited no detectable GFP signal (Fig. 6C).

**Fig. 6.**
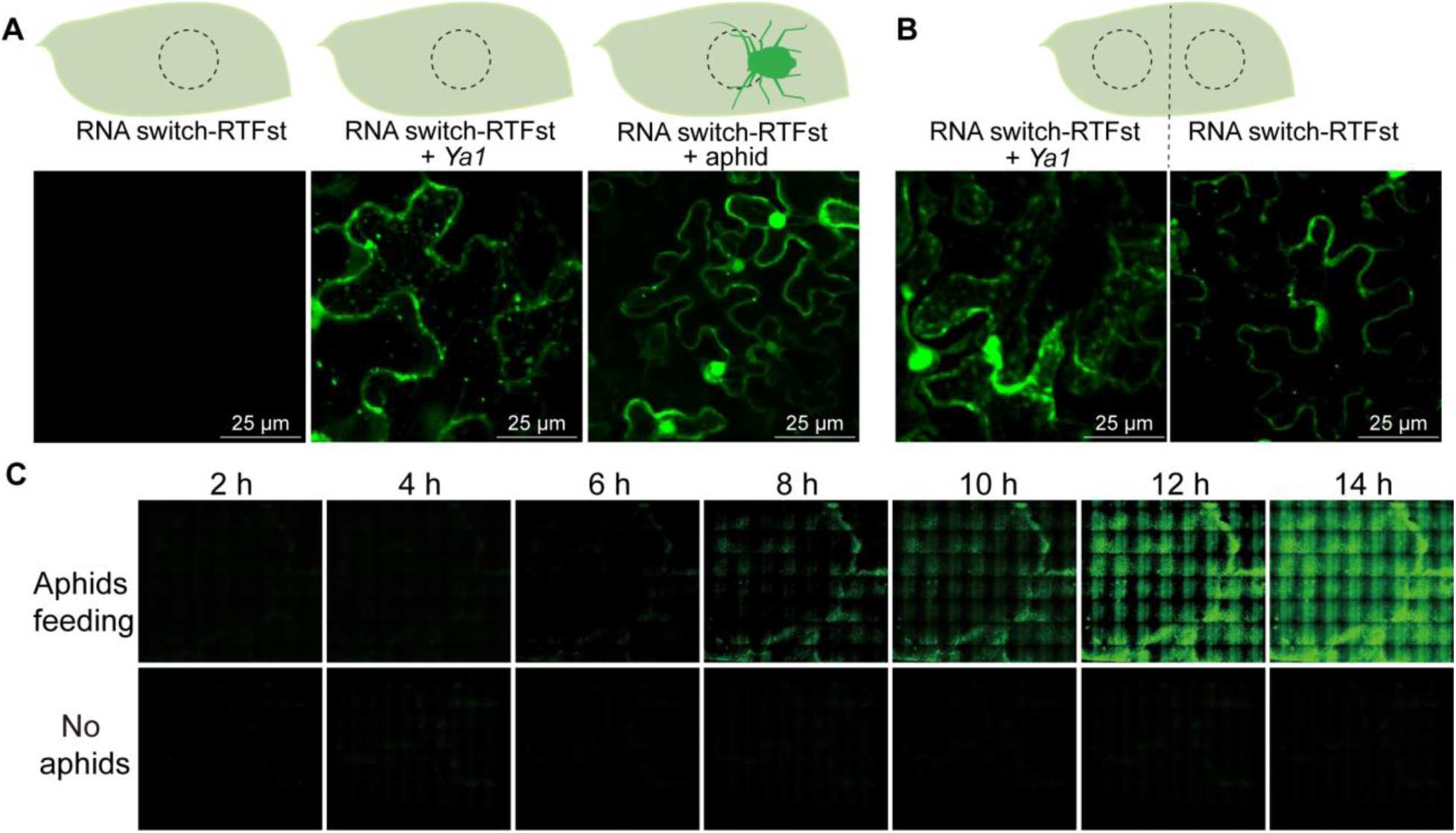
Aphid-secreted lncRNA *Ya1* is capable of cross-kingdom transfer into leaves of *N. benthamiana*. (**A**) Visualization of the aphid-secreted lncRNA Ya1 in *N. benthamiana* leaves by RNA switch– RTFst. RNA switch–RTFst + *Ya1*, RNA switch–RTFst and *Ya1* were co-expressed via agroinfiltration. RNA switch–RTFst + aphid: aphids fed on *N. benthamiana* leaves in which RNA switch–RTFst was expressed. (**B**) Visualization of cross-cell movement of *Ya1* RNA in *N. benthamiana* leaves (**C**) Visualization of the time-dependent accumulation of Aphid*-*transferred *Ya1* RNA in *N. benthamiana* leaves using RNA switch–RTFst. Images are panoramic scanning (1cm×1cm) of leaf samples using a confocal fluorescence microscope (Olympus SpinSR10). See **Video S2** for details.

These findings confirm the mobility of an aphid-secreted cross-kingdom RNA into the host plant.

## Discussion

Monitoring RNA expression dynamics in living systems at the whole-organism scale remains a fundamental challenge in biology. Here, we have addressed this issue by introducing RNA switch–RTF, a *de novo* designed tool for visualizing RNA expression dynamics in living plants. RNA switch–RTF enables high-sensitivity detection of target RNAs while preserving their native localization and function.

Live-cell RNA-imaging technologies have become indispensable for quantitatively analyzing expression dynamics^51^. However, such methods have limitations that hinder monitoring of real-time RNA dynamics at the whole-organism scale. Nucleic acid molecular beacons ^21^ and nanoparticle-based nano-flares ^52^ produce fluorescence upon structural reconfiguration triggered by RNA binding. However, efficient delivery of these exogenously synthesized probes into plant cells remains a major challenge due to the barrier of cell walls ^3^. Fluorescent RNA aptamers and RNA motifs that recruit RNA-binding fluorescent proteins offer another strategy for RNA visualization ^53–55^, but require tissue separation, which disrupts native RNA expression patterns and compromises spatial regulatory information. CRISPR–dCas-based imaging enables visualization of diverse RNAs and their subcellular localization ^56^, yet its application in plant cells is confounded by cytosolic fluorescence artifacts ^41^. A split-ribozyme system was introduced as an alternative for *in vivo* RNA imaging in plants ^28^. This approach disrupts sfGFP translation by incorporating a self-splicing ribozyme into its coding sequence, allowing translation to proceed only after the ribozyme excises and transcript ligation. While promising when target RNA and sfGFP translation occur in the same cell, the temporal delay between transcript rejoining and protein synthesis constrains its utility for real-time RNA tracking.

The effectiveness of RNA switch–RTF relies on RNA-triggered fluorescence (RTF) and programmable RNA switches. A critical component of RTF is the degron RRRG, which drives continuous GFP degradation. While several degrons have been identified in plants, including the Auxin-inducible degron (AID) ^33^ and mini-Auxin inducible degron (mIAA) ^32^, their lengths, ranging from 67–110 amino acids for AID and 40–50 amino acids for mIAA, render them unsuitable for development into RNA-binding peptides. To overcome this limitation, we conducted a proteome-wide screening of constitutively degraded proteins in plants, leading to the identification of RRRG, a 4-amino-acid degron located at the C-terminus of Niben261Chr17g0072018. This represents the smallest known degron identified in plants to date. RRRG did not disrupt the RNA-binding activity of select RBPs or their interaction with specific RNA ligands.

The second key component of RNA switch–RTF is the RNA switch itself, which regulates the degradation of GFP-RRRG. Exploiting the high affinity between Rev–RRRG and the RNA aptamer RRE ^38^, we engineered RNA switches by incorporating probe and linker sequences into RRE. Systematic testing of probe–linker combinations led to the identification of dumbbell-shaped RNA switches capable of modulating GFP degradation in response to specific RNAs. This regulation is mediated by RNA allostery, wherein binding to target RNAs induces conformational changes that alter the ability of RRE to interact with the degron on GFP-RRRG. These RNA switches represent the first known successfully engineered RNA-based regulatory elements in plants, demonstrating robust functionality across transient expression systems, including protoplasts and agroinfiltrated leaves, as well as in stable-transgenic lines.

Integration of the RNA switch–RTF system with SunTag ^40^ markedly enhances the sensitivity and precision of RNA detection by degrading free GFP. Crucially, the RNA switch–RTFst complex does not perturb the localization, expression, turnover, or translation of target RNA molecules. It also preserves target RNA function, as demonstrated by the successful compensation of shoot-derived *TCTP1* mRNA in the root tissues of *tctp1* mutants expressing the RNA switch– RTFst complex. Beyond its utility in tracking RNA dynamics in living plants, RNA switch–RTF holds promise in RNA-triggered immune responses. By engineering the system to detect pest-secreted RNAs ^10, 11^, it could be repurposed to trigger the accumulation of pest-specific defense molecules instead of GFP. Similarly, many plant pathogens secrete RNAs into host cells even at early infection stages ^50, 57, 58^. RNA switch–RTF can be designed to recognize pathogen-derived RNAs and activate species-specific immune responses. However, the system has several limitations. The RNA switch requires a stable dumbbell-shaped structure, constraining its compatibility to transcripts with suitable sequences. Additionally, the ∼21-nt probe in the RNA switch may exhibit off-target effects. Probe accessibility depends on maintaining an unpaired configuration, which can be disrupted by intracellular or extracellular factors that alter RNA structure, compromising target recognition and overall switch performance.

These findings establish RNA switch–RTF as a versatile platform for RNA imaging and functional studies while opening new and sustainable avenues for enhancing plant immunity against pests and pathogens.

## Materials and Methods

### Plant materials and growth conditions

Seeds of *Nicotiana benthamiana* and *A. thaliana* Columbia (Col-0) were surface-sterilized and germinated on 1× Murashige and Skoog (MS) medium with 1% [w/v] Sucrose. For normal growth conditions, plants were grown at 23°C under long-day conditions (LD, 16-h light /8-h dark cycle) with a light intensity of 150 μmol m^-2^ s^-1^ in a plant growth chamber.

The *A. thaliana* mutants *cm3* (SALK_011276.36.40.x) *and tctp1* (CS486456, heterozygous) were obtained from the Arabidopsis Biological Resource Center (ABRC). Homozygous plants were used and primers for genotyping are listed in Table S7.

For tobacco *mCherry* lines, we introduced *35S::mChery* construct into *Nicotiana benthamiana* by Agrobacterium tumefaciens-mediated transformation. The overexpression lines were confirmed by quantitative real-time PCR (qRT-PCR).

### Agroinfiltration of *N. benthamiana*

*Agrobacterium tumefaciens* GV3101 stocks were streaked on LB plates supplemented with kanamycin (50 μg/mL) and rifampicin (25 μg/mL). After overnight culture in liquid LB, cells were pelleted, resuspended in infiltration medium (10 mM MES, pH 5.6; 10 mM MgCl₂; 150 μM acetosyringone), and adjusted to an OD_600nm_ of ∼0.6. After a 2-h incubation, the suspension was infiltrated into the abaxial side of 3-week-old *N. benthamiana* leaves using a needleless syringe. Co-infiltration used a 1:1 volume ratio of bacterial solutions containing constructs of interest. Plants were maintained in growth chambers (LD; 16-h light /8-h dark cycle, with a light intensity of 150 μmol m^-2^ s^-1^) for 3 d before leaf samples were collected for confocal microscopy.

### Identification of putative degrons in *N. benthamiana*

Leaves of 6-week-old *N. benthamiana* were treated with either 10 μM MG132 or mock solution (1× PBS buffer) for 24 h, respectively. Following the treatment, leaves were harvested for LC–MS/MS at Shanghai Personalbio Technology Company. Proteins were extracted using TCA/acetone precipitation and the SDT (4% SDS, 100 mM DTT, 150 mM Tris-HCl pH 8.0) lysis method^59^. The mixture was sonicated, boiled, and centrifuged to separate the supernatant. Protein concentration was determined using the BCA Protein Assay Kit (Cat. A65453, Thermo Scientific), and 15 µg of protein from each sample was mixed with 5× SDS loading buffer and boiled. The proteins were separated on a 4–20% SDS–PAGE gel and visualized with Coomassie Blue R-250 staining. Protein digestion was performed by adding DTT (40 mM) to each sample, mixing at 600 rpm for 1.5 h (37℃), and then adding IAA (20 mM) to block reduced cysteine residues. The samples were then transferred to filters (Microcon units, 10 kDa), washed with UA buffer (8 M urea, 100 mM Tris-HCl pH 8.0) and 25 NH_4_HCO_3_ buffer, then trypsin was added (trypsin:protein (w/w) ratio was 1:50) and incubated at 37℃ for 15–18 h. The resulting peptides were collected, desalted on C18 Cartridges (Cat.66871-U, Sigma), concentrated by vacuum centrifugation, and reconstituted in 0.1% v/v formic acid.

For the analysis of data-independent acquisition (DIA) experiments, iRT (indexed retention time) calibration peptides were spiked into the sample and analyzed by an OrbitrapTM AstralTM mass spectrometer connected to a Vanquish Neo system liquid chromatography in DIA mode. The precursor ions were scanned at a mass range of 380–980m/z, and the MS1 resolution was 240,000 at 200 m/z. DIA data were analyzed with DIA-NN software ^60^, and protein identification was based on 99% confidence and a false-discovery rate ≤ 1%. Proteins were identified by searching MS/MS against the InterPro member database Pfam.

To evaluate the degradation efficiency of putative degron sequences, the 60 nucleotides (corresponding to 20 amino acids) for each gene were fused to the C terminus of the GFP sequence in a 2A self-cleaving peptide co-expression plasmid (Table S1). This plasmid also co-expresses mCherry alongside GFP. Various truncated versions of CP-12 (Niben261Chr17g0072018) were generated and inserted into the plasmid. These plasmids were introduced to *N. benthamiana* leaves via agroinfiltration. Degradation efficiency was assessed by comparing GFP signal intensity to that of mCherry. Fluorescence images were captured using a Leica SP5 X MP confocal microscope using a 40x Leica objective, with excitation and emission at 488 and 509 nm for GFP, and 575 and 650 nm for mCherry.

### Synthesis of RNA-binding peptides, RNA aptamers, and electrophoretic mobility shift assay (EMSA)

The RNA-binding peptides (Rev–RRRG, P22N–RRRG, λN–RRRG, BTat–RRRG, HTat– RRRG, and Rex–RRRG, Table S2) were synthesized using standard Fmoc solid-phase synthesis at Sangon Biotechnology Company. Briefly, the synthesis of peptides was carried out on ChemMatrix resins using Rink Amide to produce C-terminal amides. For the coupling of amino acids, a pre-activation step was conducted using five equivalents of Fmoc-protected amino acid with 4.9 equivalents of *O*-(benzotriazol-1-yl)-*N*,*N*,*N*′,*N*′-tetramethyluronium hexafluorophosphate (HBTU), five equivalents of N-hydroxybenzotriazole (HOBt), and 10 equivalents of *N*,*N*-diisopropylethylamine (DIPEA) in dimethylformamide (DMF), followed by reaction with the N-terminally deprotected peptidyl resin. Thorough washing of the resin with DMF was executed, and the completion of the reaction was assessed using the Kaiser test. Deprotection of Fmoc was achieved with an excess of 20% (v/v) piperidine in DMF, and the washed deprotected peptidyl resin was rinsed with DMF to eliminate any residual piperidine. Cleavage from the resin along with side-chain deprotection occurred using a mixture of 95% TFA, 2.5% TIS, and 2.5% H_2_O at room temperature for 1.5–2 h. The resultant peptide was precipitated using cold diethyl ether, separated via centrifugation, and subsequently dissolved in water containing 0.1% v/v trifluoroacetic acid before undergoing RP–HPLC and ESI–MS analyses.

The reverse complementary sequences of RNA aptamers’ DNA (RRE, P22N, λN, Pepper, and XBE (Table S8) with *T7* promoter were synthesized and annealed with the *T7* promoter as templates for *in vitro* transcription. The RNA aptamers were *in vitro* transcribed according to the Transcription T7 kit (Cat. 6140, Takara) and labeled with biotin using Pierce™ RNA Biotin Labelling Kit (Cat. 20160, Thermo Scientific).

EMSA experiments were performed using a LightShift Chemiluminescent RNA EMSA Kit (Cat. 20158, Thermo Scientific). 3′ biotin-labeled RNA probes and synthesized peptides were incubated in binding buffer (3 mM MgCl_2_, 40 mM KCl, 5% v/v glycerol, 2 mM DTT, 2 μg tRNA) for 40 min at room temperature, and mobility shift assay was performed using 6% acrylamide native gel electrophoresis. After electrophoresis, gels were sandwiched between filter paper and 0.5x TBE buffer-soaked nylon membrane in a clean electrophoretic transfer unit for electrophoretic transfer at 400 mA (∼35 V) for 30 min. After UV cross-linking, biotin signals were detected with stabilized streptavidin–horseradish peroxidase conjugate using a Chemiluminescent Nucleic Acid Detection Module (Thermo Fisher Scientific, China).

### Design of RNA switches

RNA switches were designed based on the RRE sequence, containing three functional elements, probe sequence, linker 1, and linker 2. Linkers 1 and 2 hybridize with the RRE and the probe, respectively.

Linker 2 is designed as the reverse complementary sequence to the 3′ end of the RRE ligand. Its role is to disrupt the active conformation of the RRE ligand by base-pairing with its 3′ end. Therefore, the linker 2 sequence should be placed at the 5′ end of the RNA switch, allowing it to hybridize with the 5′ end of the RRE ligand, which is critical for maintaining the inactive conformation of the RNA switch. The binding strength between Linker 2 and the RRE ligand should not be excessively strong. If the binding force is too strong and the change of RRE conformation is completely disrupted, the RRE will not be able to revert to its active conformation, even when the probe binds.

Linker 1 sequences were introduced, which is designed to be reverse complementary to the 3′ end of the probe sequence. Its primary function is to enhance the inactive conformation of the RRE by Linker 2 in the absence of the target RNA. In the presence of the target RNA, the probe binds to the target RNA, triggering the dissociation of linker 1 from the probe. This dissociation further disrupts the binding of linker 2 and the RRE ligand, allowing the RRE to regain active conformation.

For probe design, we used Stellaris Probe Designer on Biosearch Technologies (https://www.biosearchtech.com/support/tools/design-software/stellaris-probe-designer) with default settings. The 5′ and 3 UTR regions of target RNAs should be avoided in designing the probe. A probe with an annealing temperature below 65°C and a minimum length of 22 nucleotides is desirable.

The structures of RNA switch sequences were predicted by mFold (www.unafold.org/mfold/applications/rna-folding-form.php) with default settings.

### Expression of RNA switch–RTF in plants

The plasmid pCambia1300 was used as a backbone to construct RNA switch-RTF components. The sequences encoding fused GFP–RBP–RRRG proteins were cloned into the plasmid under the *UBQ10* promoter. The sequences for the RNA switch driven by the *AtU6* promoter, with various probes targeting *mCherry*, were cloned in the same plasmid upstream of the *UBQ10* promoter. The RNA switch sequences are listed in Table S9. These types of plasmids were named RNA switch– RTF plasmids (Table S10) and were introduced into the leaves of transgenic *N. benthamiana mCherry* lines via agroinfiltration.

The RNA switch–RTF plasmid with a probe targeting *CM3* mRNA was introduced into *A. thaliana* wild type and *cm3* mutant plants respectively via floral dip. Homozygous T_3_ lines were used.

### Design of RTF–SFP and RTF–SunTag

RTF–SFP was modified in the RNA switch–RTF plasmid described above by replacing sequences encoding fused GFP–Rev–RRRG proteins with the sequences of GFP_1–10_ (the GFP β-strand 1-10, which could combine the GFP β-strand 11 to restore GFP signal), an internal ribosome entry site (IRES), and 24× GFP_11_. These were named RNA switch–RTF–SFP plasmids (Table S10).

RTF–SunTag was also modified in the RNA switch–RTF plasmid, by replacing sequences encoding fused GFP–Rev–RRRG proteins with sequences of SunTag–Rev, an internal ribosome entry site (IRES), and GFP–scFv–RRRG. These were named RNA switch–RTF–SunTag plasmids (Table S10).

### Generation of transgenic *A. thaliana* plants expressing various RNA switch–RTFst constructs

A series of RNA switch–RTF–SunTag (RNA switch–RTFst) plasmids with various probe sequences in RNA switches were constructed (Table S9). These were introduced to *A. tumefaciens* GV3101 and transgenic *A. thaliana* lines were generated via the floral-dip method *(2)*, and homozygous lines were used.

### Single-molecule fluorescence *in situ* hybridization

For probe design, five probes targeting various sites of the *CM3* coding region were designed using Stellaris® Probe Designer version 2.0 from Biosearch Technologies (http://singlemoleculefish.com). Before ordering pre-labeled probes from Biosearch Technologies, a TAIR BLAST query was run for each sequence to ensure target specificity (https://www.arabidopsis.org/Blast/). The oligonucleotides of each probe, labeled with Cy3 at the 5′ terminus, were synthesized by Sangon Biotech Co., Ltd..

For sample preparation, root tips of 10-d-old transgenic seedlings were collected and fixed in 2-(N-morpholino) ethane sulfonic acid (MES) buffer, pH 5.7, containing 4% v/v paraformaldehyde for 30 min at room temperature. The roots were rinsed twice with 1× PBS and placed onto a microscope slide with a coverslip, gently squashed and briefly submerged in liquid nitrogen. The coverslips were then removed, and the samples were left to dry at room temperature for 30 min. Tissue permeabilization was carried out by immersing the samples in Coplin jars containing 70% v/v ethanol, which were shaken gently for at least 1 h. The ethanol was evaporated at room temperature before the roots were washed twice with MES.

Hybridization was performed as previously described (*3*). Briefly, 100 μL of hybridization solution with probes at a final concentration of 250 nM was added to each slide. Coverslips were placed over the samples to prevent buffer evaporation, and the probes were left to hybridize in a humid chamber at 37°C overnight in the dark. The samples were washed twice for 45 min at 37°C with freshly prepared 1× SSC/40% v/v formamide buffer, rinsed at room temperature with MES buffer (pH 5.8), and dried. A drop of ProLong Diamond Antifade Mountant (Invitrogen) containing DAPI was added prior to observation.

### Construction of MCP–MS2 system and dCas9-SunTag system

The DNA sequence of 24× MS2 was cloned into the 3′ UTR of *CM3* under control of the *UBQ10* promoter to form a fused CM3–24×MS2 expression plasmid. The DNA encoding fused mCherry–MCP was cloned into the plasmid under the *UBQ10* promoter. The DNA of <u>dCas9-SunTag were synthesized and subsequently inserted downstream the</u> *UBQ10* promoter in a pCAMBIA1300s plasmids containing a cognate sgRNA expression cassette.

### Protoplast isolation and imaging

*Agrobacteria* containing RNA switch–RTF–SunTag plasmids were co-transformed into tobacco epidermal cells with containing CM3–24×MS2 plasmids and the mCherry–MCP plasmid by agroinfiltration as described above. Twenty-eight hours after infiltration, leaves were cut off for protoplast isolation based on a described previously protocol ^61^. Images were taken using a Leica SP5 X MP confocal microscope using a 40x Leica objective, with excitation and emission at 488 and 509 nm for GFP, and 575 and 650 nm for mCherry.

### Estimating the proximity of RNAs to mitochondria

*Nicotiana benthamiana* leaves were agroinfiltrated with constructs expressing RNA switch-RTF-SunTag and mitochondrial-targeting RNA *VDAC3*. Twenty-eight hours after infiltration, leaves were infiltrated with 1 μM MitoTracker® Red CMXRos (ThermoFisher, A66443) for mitochondria labeling. The movements of *VDAC3* RNA particles to mitochondria in living epidermal cells were captured using confocal microscopy time-lapse imaging. The distance from the center of each *VDAC3* RNA particle to the center of the closest mitochondrion is measured with ImageJ and the number of particles at a given mitochondrion distance is calculated ^44^.

### Western blotting

Total protein extraction and western blotting were performed as previously described (*5*). Briefly, protein extracts were mixed with SDS–PAGE sample buffer and denatured at 95°C for 5 min. After brief centrifugation, denatured protein samples were separated by 12% SDS–PAGE and blotted onto PVDF membranes. Primary and secondary antibodies were used at the following dilutions: 1:10,000 for GFP (ab290; Abcam), 1:10,000 for ubiquitin (MAB3408; Sigma-Aldrich), and 1:10,000 for anti-IgG (mouse) HRP conjugate (W402B; Promega). Immunoreactive bands were visualized using ImmunoStar® LD (Wako).

### Microscopy setup and image acquisition

For protoplast imaging, protoplasts were stained with 100 µM FM4-64 for 1 min and washed with W5 solution before imaging. For epidermal-cell imaging, agroinfiltrated *N. benthamiana* leaves were cut for imaging 72 h after agroinfiltration. For mitochondrial localization, agroinfiltrated leaves were incubated with 200 nM MitoTracker red CMXRos in 1× PBS buffer with 3% w/v sucrose for 15 min before imaging (*6*). Images were captured on a spinning-disk confocal microscope (Olympus SpinSR10) with an EMCCD camera and 40× oil objective. GFP, Cy3, DAPI, FM4-64 and MitoTracker red CMXRos signals were detected at respective excitation/emission wavelengths of 488/498–559 nm, 554/568 nm, 405/420–480 nm, 488/734 nm and 561 nm/580–625 nm. Panoramic images of tissues were acquired with the Olympus VS200 panoramic digital scanner (Japan). For visualization of *TUB5* expression during plant growth, GFP signal in whole plants was acquired using NightShade LB 985 In vivo Plant Imaging System (Berthold) every two weeks throughout the growth period. For detection of *TCTP1* mRNA long-distance movement, Arabidopsis rootstock samples at different distances from the graft junction were cut for Z-stack image acquisition using a spinning-disk confocal microscope (Olympus SpinSR10) with an EMCCD camera and 20× objective. For observation of the cell-to-cell movement of *TCTP1* mRNA, time-lapse imaging of elongation-region cells of rootstock roots was performed using a SpinSR10 microscope with a 20× objective every 3 sec for 1 min.

### RT–qPCR

Total RNA was extracted using the Qiagen RNeasy kit, following the manufacturer’s instructions. Reverse transcription was performed with 1 μg RNA using the Qiagen Omniscript kit with a primer mix of random 10 mers (final concentration of 10 μM) and 15 mers oligo dT primers (final concentration of 1 μM). A negative control was performed by adding water instead of reverse transcriptase. For gene expression profiling, qPCR was performed with gene-specific qPCR primers. Ct readouts of each gene were first normalized to the housekeeping gene *β-Actin* (ΔCt), and the relative expression of individual genes versus the expression levels in control conditions was then calculated with the 2^−ΔΔCt^ method. Primers are listed in Table S7.

### Micro-grafting of *A. thaliana* plants

*Arabidopsis thaliana* seeds were germinated and grown vertically on plates containing 1/2 MS salts, 1% w/v sucrose, and 1% w/v micro agar under short-day conditions in a controlled-environment chamber (Percival) for one week. Micro-grafting of seedlings was performed as described (*7*). Briefly, seedlings of similar sizes and with evenly elongated hypocotyls were cut in the upper half of the hypocotyl with a sterilized razor blade. The junctions of the graft were supported using silicon microtubing (0.3-mm internal diameter). Grafted plants were then transferred to new plates and grown vertically under short-day conditions. Adventitious roots that formed above the junction were removed every 2 d.

### Aphid feeding experiments

*Ya1*-targeted RNA switch–RTFst plasmids were agroinfiltrated into four-week-old *N. benthamiana* leaves. Forty-eight hours post-infiltration, five aphids were confined to the infiltration zone using rubber bands and double-sided tape to prevent escape. After 24 h of feeding, leaf samples were collected for confocal fluorescence microscopy using an Olympus FV3000. Additionally, co-infiltration of full-length *Ya1* plasmids with *Ya1*-targeted RNA switch-RTFst plasmids was performed.

Leaf discs (4 cm²) containing agroinfiltrated switch–RTFst plasmid constructs were excised 48 h post-infiltration and transferred to six-well plates containing Murashige and Skoog (MS) liquid medium to maintain osmotic balance and humidity. Aphids (3–4 individuals) were introduced onto the leaf discs, followed by incubation in controlled growth chambers (25°C, 16/8 h light/dark). Time-lapse imaging was conducted at hourly intervals using an Olympus SpinSR10 spinning-disk confocal microscope, equipped with an automated stage scanning system for panoramic imaging.

## Data availability

All data are available in the main text or the supplementary materials. All materials (seed stocks, plasmids) are available upon request.

## Acknowledgments

We sincerely thank Professor Yuelin Zhang (Sichuan University) for his insightful discussions and critical reading of the manuscript. We also appreciate Dr. Yiliang Ding (John Innes Centre, UK) and Dr. Xiaofei Yang (Center for Excellence in Molecular Plant Sciences, Chinese Academy of Sciences, China) for their valuable suggestions.

## Author contributions

Conceptualization: JB, YZ, YC, CF

Methodology: JL, SP, YT, ML

Investigation: JB, JZ, HR

Visualization: SL, JY, MD, FC

Funding acquisition: JB, YZ, YC, JL

Project administration: YZ, CF

Supervision: YZ, CF

Writing – original draft: JB

Writing – review & editing: YC, YZ, CF

## Supplementary Materials for

The PDF file includes:

Figs. S1 to S5

Tables S1 to S10

References (1-13)

**Other Supplementary Materials for this manuscript include the following:**

Video S1

Video S2

---

**Fig. S1.**
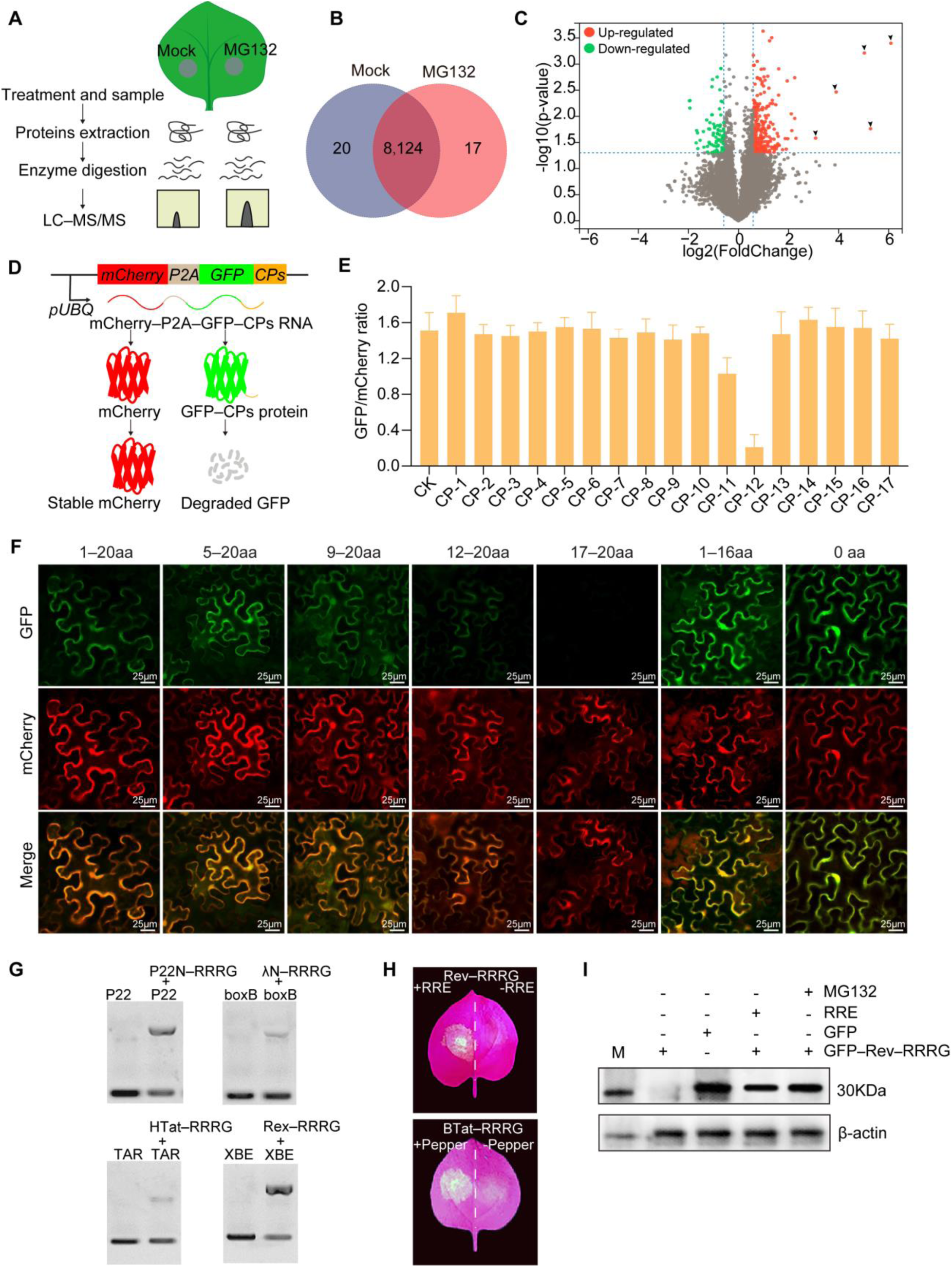
*De novo* identification of protein degron sequences in *N. benthamiana*. (**A**) Flowchart showing the identification of constitutively degraded proteins. *Nicotiana benthamiana* epidermal cells were infiltrated wither with 10 μM MG132 for 24 h to inhibit proteasome activity or 1× PBS buffer (mock), followed by protein extraction and LC–MS/MS analysis. (**B**) Seventeen proteins were identified in *N. benthamiana* leaves treated with MG132 compared to those treated with the mock groups. (**C**) Volcano plot illustrating differentially abundant proteins. (**D**) Schematic of the co-expression system used for fluorescence normalization of GFP fusions. mCherry and GFP, tagged with the putative degron sequence, were co-expressed using a 2A self-cleaving peptide system. CP represents the 20 aa at the C-terminal of 17 proteins detected in MG132-treated leaves. (**E**) Assessment of the effect of CPs on GFP degradation based on the GFP/mCherry fluorescence intensity ratio. CK means control check. (**F**) Representative image showing the effect of different truncations of CP-12 on GFP degradation. Scale bar = 20 μm. (**G**) Gel-shift assay testing RNA-binding ability of RBP tagged with the RRRG degron. (**H**) GFP fluorescence signals in the presence and absence of specific RNA aptamers when BTat and Rev are tagged with RRRG. *N. benthamiana* leaves were imaged at 3 d post-agroinfiltration. (**I**) Western blot detection of GFP.

**Fig. S2.**
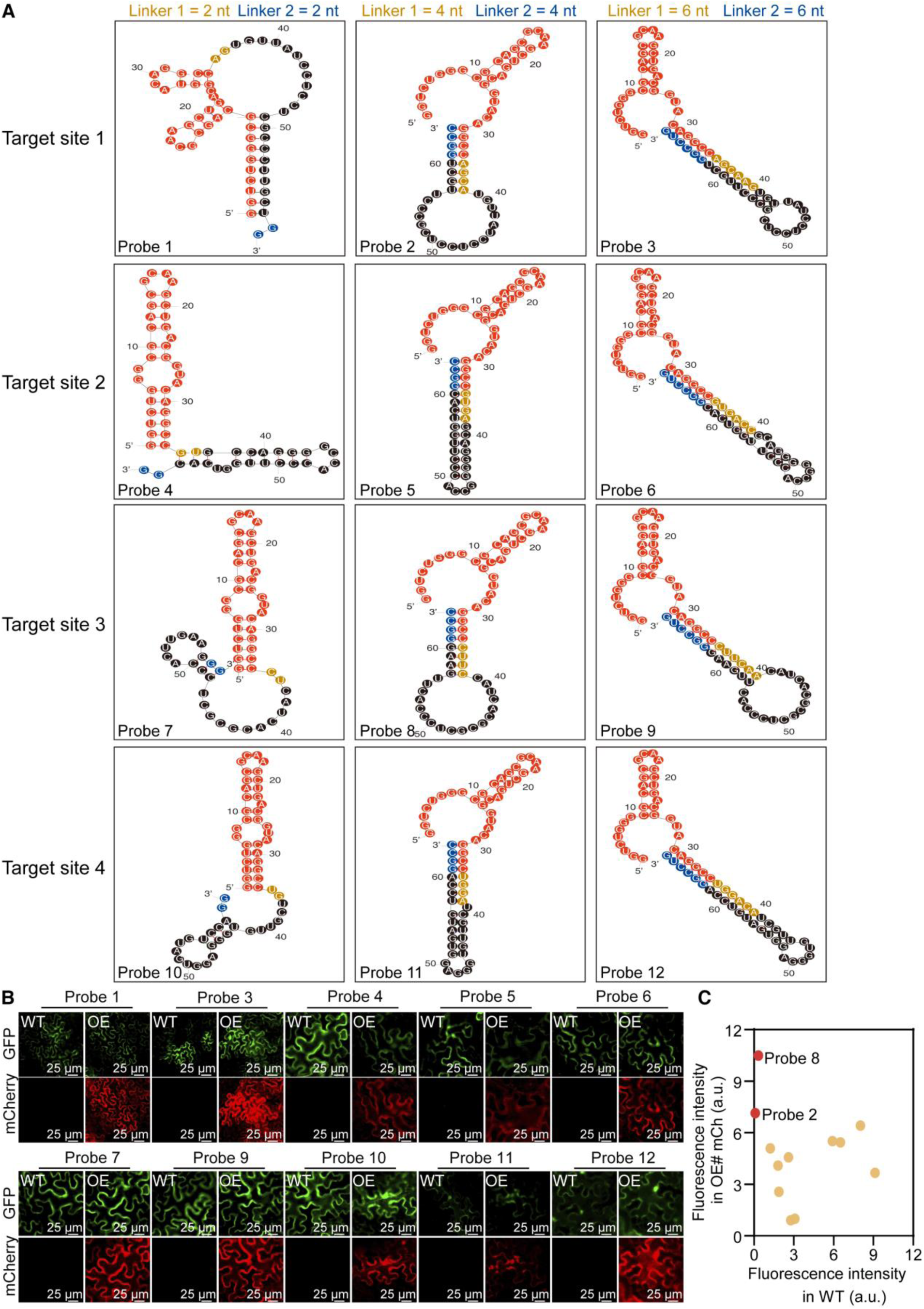
Designing of RNA switches targeting *mCherry* RNA. (**A**) Minimal free energy (MFE) secondary structure of 12 RNA switches (Probes 1–12) designed for specific targeting of *mCherry*. The RNA switches feature linker lengths of 2–6 nt. (**B**) Screening of RNA switches using fluorescent microscopic images of *N. benthamiana* epidermal cells. Scale bars = 50 μm. (**C**) Fluorescence quantification of 12 RNA switches to control RTF reporter expression in WT and *mCherry*-expressing epidermal cells of *N. benthamiana* leaves.

**Fig. S3.**
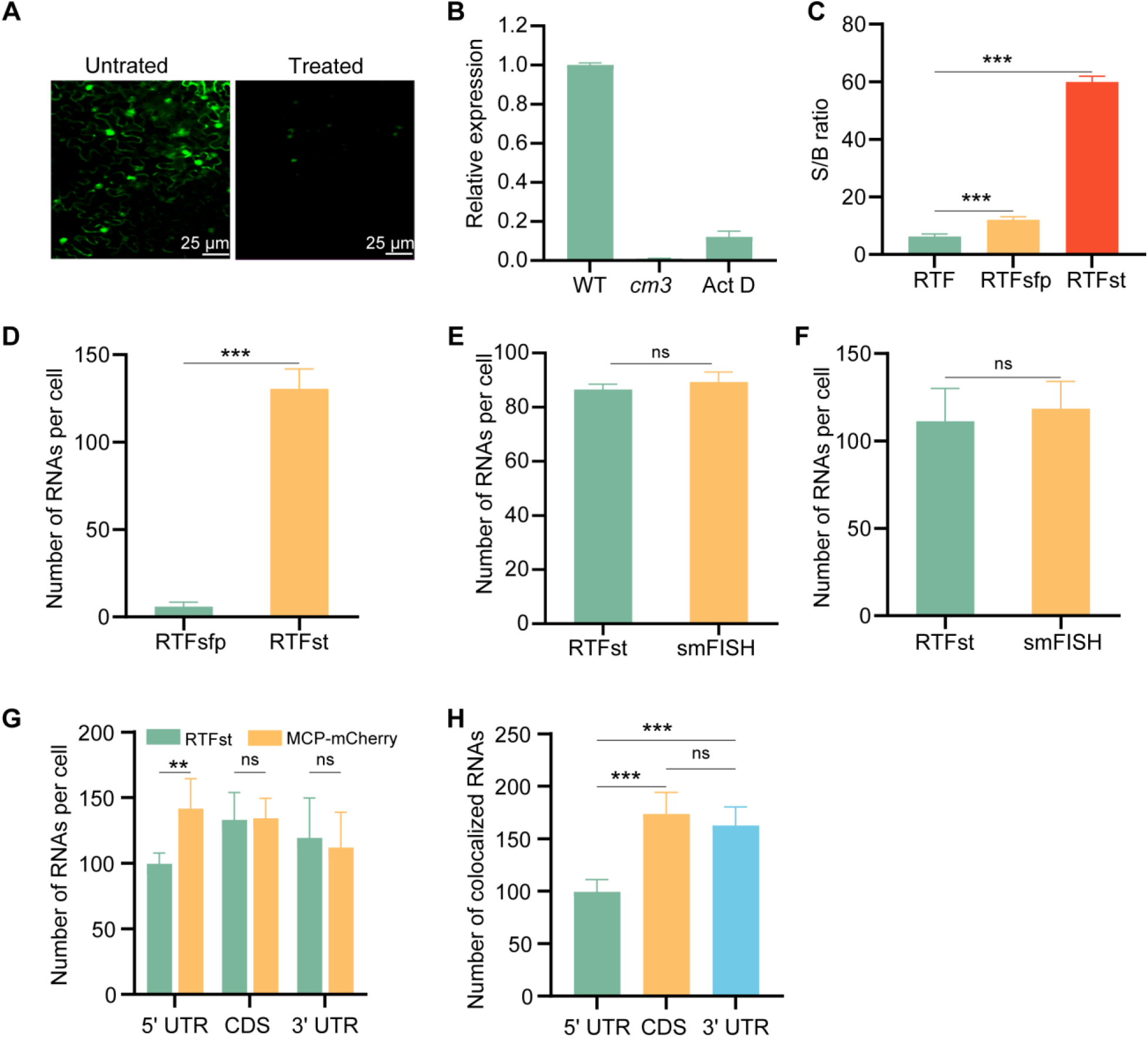
Expression of RNA switch-controlled RTF system in plants. (**A** and **B**) The RNA switch–RTF system exhibits a degree of leakage in the absence of the target RNA. (**A**) Fluorescence imaging of *N. benthamiana* leaf epidermal cells expressing the RNA switch–RTF system targeting *CM3* mRNA. ‘Untreated’ indicates cells not exposed to Actinomycin D (Act D), an RNA synthesis inhibitor, while ‘treated’ indicates cells subjected to 20 μM Act D treatment. Scale bars: 25 μm. (**B**) Relative expression levels of *CM3* mRNA in wild-type (WT), *cm3* mutant, and Act D-treated WT plants determined by qRT–PCR. (**C**) Quantification of the signal-to-noise ratio (SNR) for RTF, RTF–SFP, and RTF–SunTag in *cm3* mutants (n = 6, mean ± s.e., *** *P* < 0.001, one-way ANOVA with multiple comparisons and a post-hoc test). (**D**) The number of well-dispersed diffraction-limited fluorescent puncta per cell (n = 6, mean ± s.e.m., **** *P* < 0.001, two-tailed unpaired Student’s t-test). **(E** and **F)** Quantification of *CM3* mRNA in individual root (**E**) and leaf (**F**) cells using smFISH and RNA switch–RTFst. Statistical comparisons for root cells (n = 6, mean ± s.e., *P* > 0.05, non-significant [ns], two-tailed unpaired Student’s t-test) and leaf cells (n = 6, mean ± s.e., *P* > 0.05, ns, two-tailed unpaired Student’s t-test). (**G**). Quantification of *CM3* mRNA detected by RNA switch–RTFst and mCherry–MCP– MS2 (n = 6, mean ± s.e, ns, *P* > 0.05; P < 0.01, two-tailed unpaired Student’s t-test). (**H**) Number of colocalized *CM3* mRNAs detected by RNA switch–RTFst and mCherry–MCP–MS2 (n = 6, mean ± s.e., \*\*\**P* < 0.001, one-way ANOVA with multiple comparisons and a post-hoc test).

**Fig. S4.**
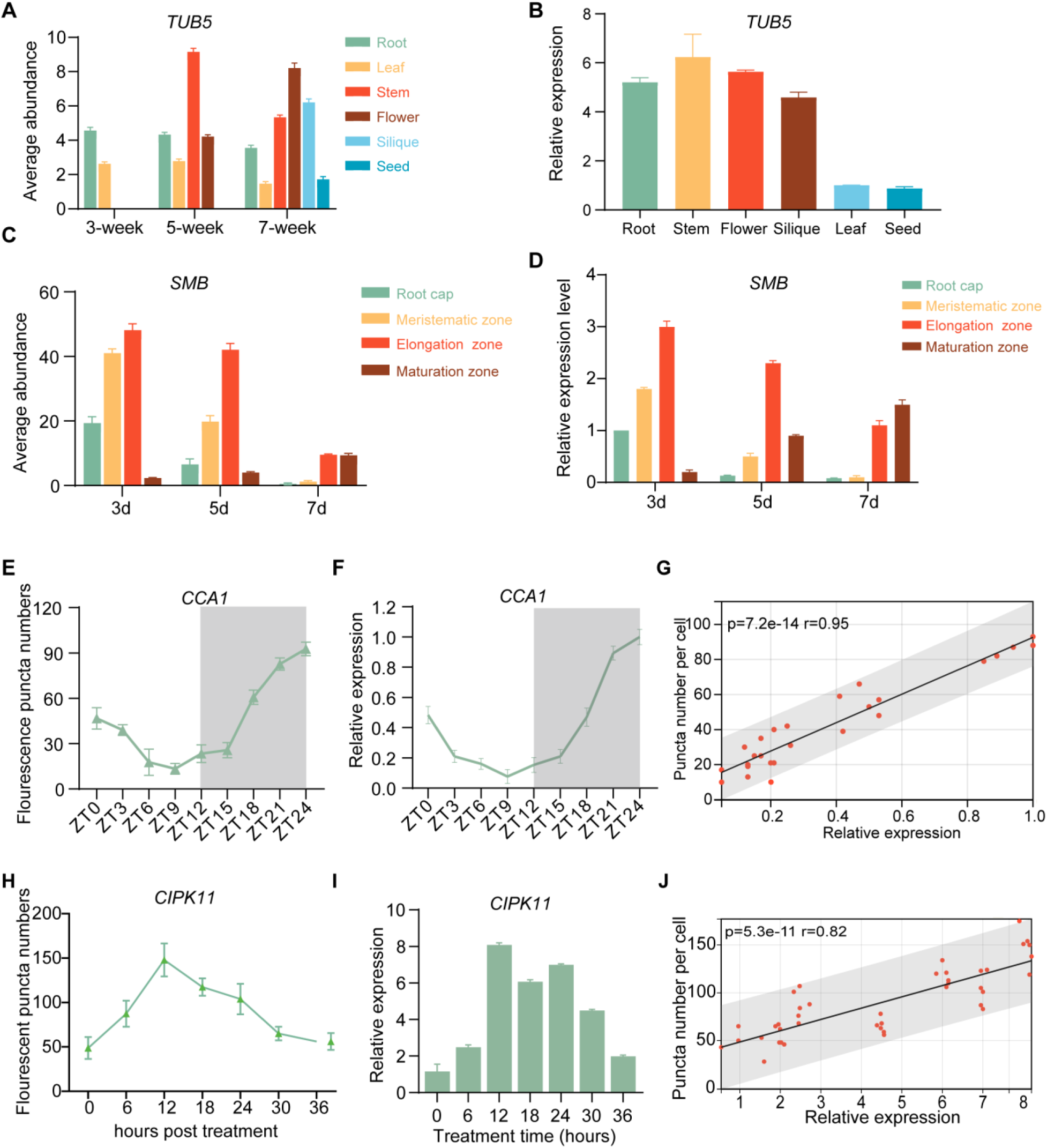
Quantification of RNA abundance using RNA switch–RTFst in living plants. (**A**) Average abundance of *TUB5* in different tissues as measured by RNA switch–RTFst. (**B**) Relative expression levels of *TUB5* in different tissues of 7-week-old *A. thaliana* seedlings assessed by RT–qPCR. (**C**) Average abundance of *SMB* in root tip tissues, as revealed by RNA switch–RTFst. (**D**) Relative expression levels of *SMB* in root tip tissues assessed by RT–qPCR at 3, 5, and 7 d after germination. (**E** and **F**) Circadian patterns of *CCA1* expression in leaves of 7-week-old *A. thaliana* depicted by RNA switch–RTFst quantification (**E**) and RT–qPCR (**F**). Light phase began at Zeitgeber Time 0 (ZT0) and dark phase (shaded areas) began at ZT12. (**G**) Correlation between *CCA1* expression levels measured by RNA switch–RTFst and RT–qPCR. (**H** and **I**) Average abundance of *CIPK11* in roots of 7-week-old Arabidopsis under cadmium (Cd) stress as determined by RNA switch–RTFst (**H**) and RT–qPCR (**I**). (**J**) Correlation between *CIPK11* expression levels measured by RNA switch–RTFst and RT–qPCR.

**Fig. S5.**
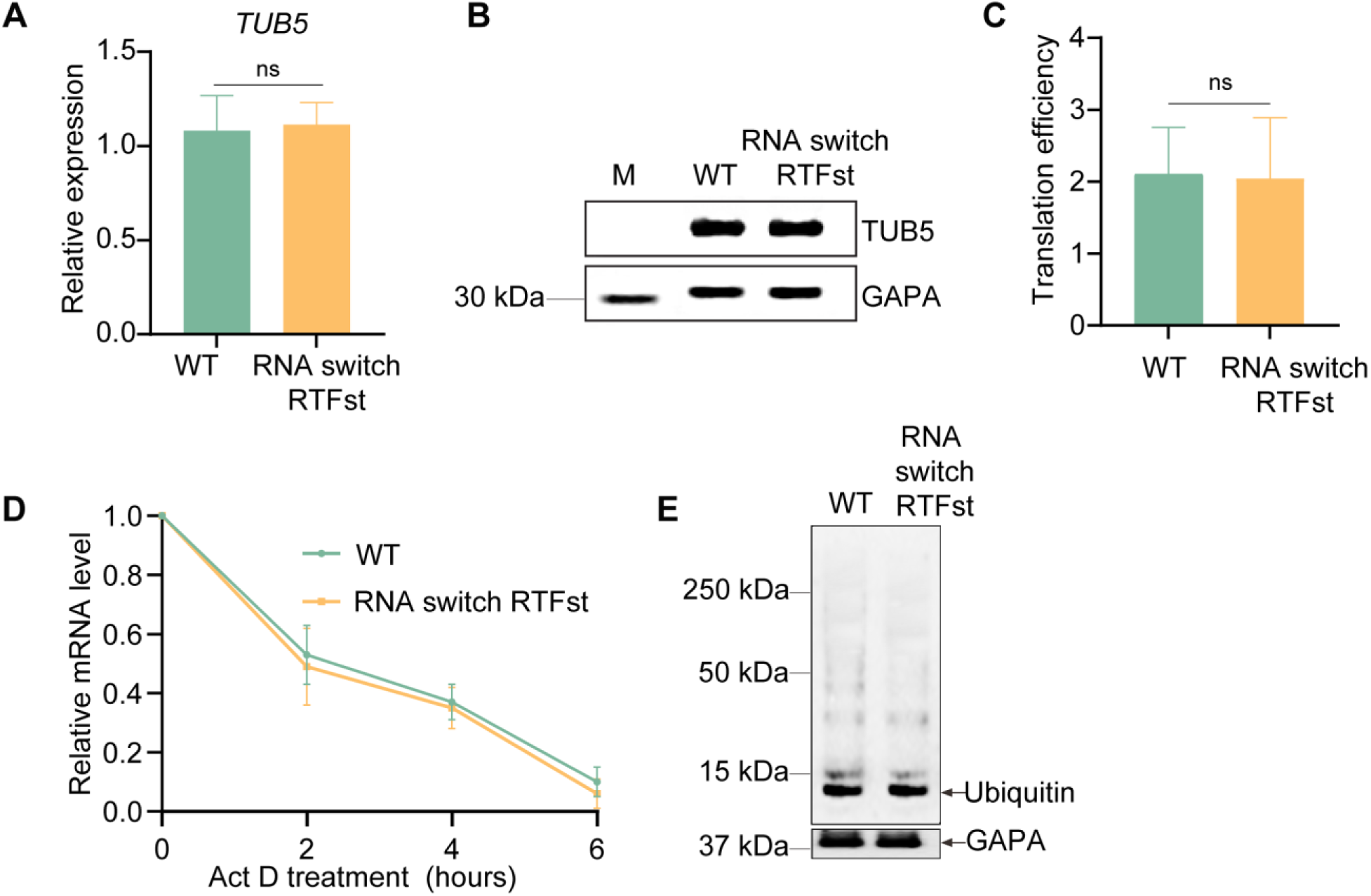
The fate of *TUB5* mRNA in Arabidopsis thaliana is unaffected by the RNA switch-RTFst system. (**A**) Relative expression levels of *TUB5* mRNA in WT plants and plants expressing RNA switch– RTFst (n = 9, mean ± s.e., ns, non-significant, *P* > 0.05, two-tailed unpaired Student’s t-test). (**B**) Western blot of TUB5 abundance in WT plants and plants expressing RNA switch–RTFst. GLYCERALDEHYDE 3-PHOSPHATE DEHYDROGENASE A SUBUNIT (GAPA) was used as a loading control. (**C**) Translation efficiency of *TUB5* mRNA in WT plants and plants expressing RNA switch–RTFst calculated by normalizing TUB5 abundance to *TUB5* mRNA levels (n = 9, mean ± s.e., ns, non-significant, *P* > 0.05, two-tailed unpaired Student’s t-test). (**D**) Degradation dynamics of *TUB5* transcripts following Act D treatment in the roots of WT and plants expressing RNA switch–RTFst. One-week-old *A. thaliana* seedlings were treated with 20 μM Act D, and relative *TUB5* transcript levels were quantified by RT–qPCR at 0, 2, 4, and 6 h post-treatment. (**E**) Western blot of the overall ubiquitinated protein in WT and switch–RTFst-expressing lines. GAPA was used as a loading control.

**Table S1.**
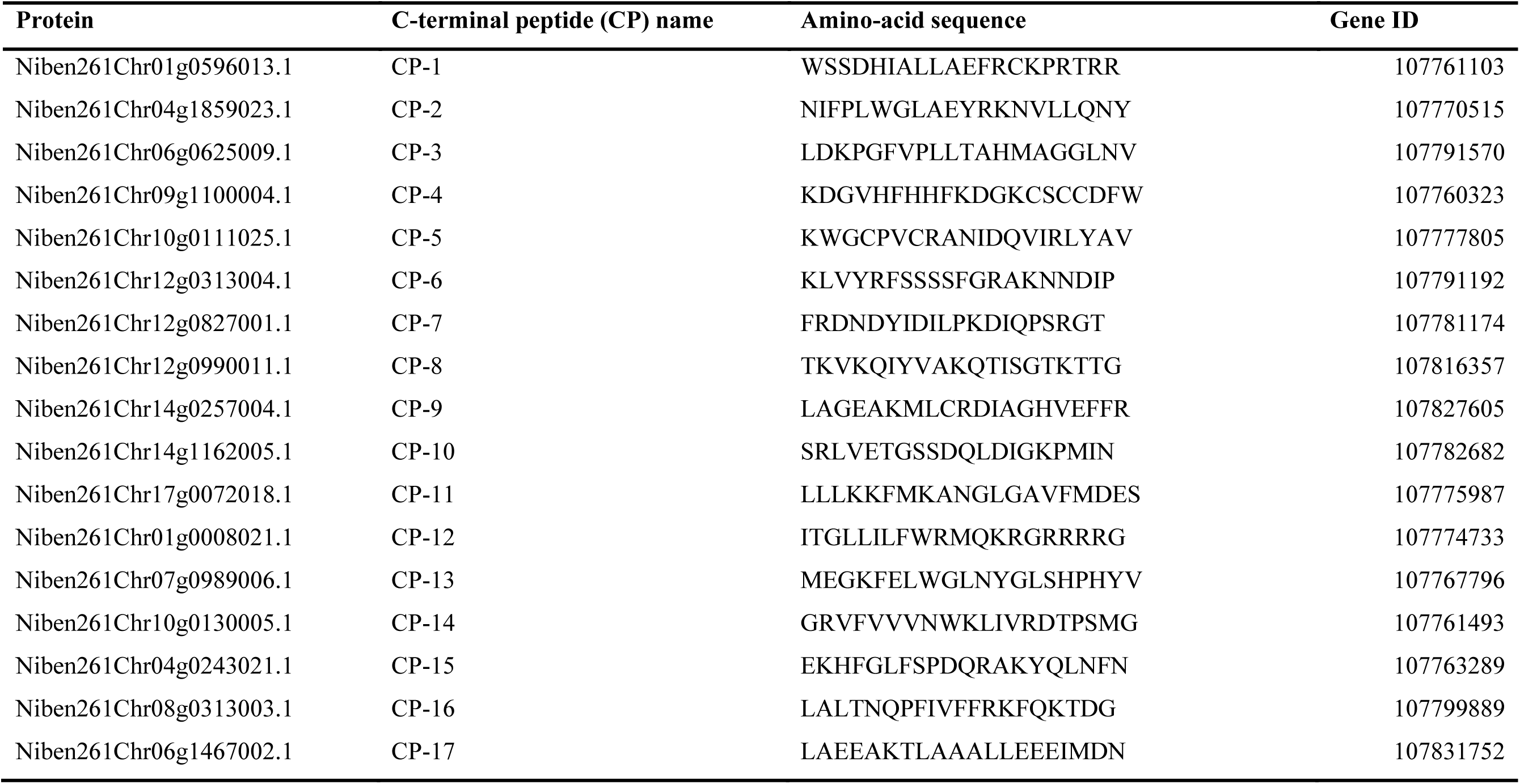
Twenty amino acids at the C-terminal of 17 proteins detected in MG132-treated *N. benthamiana* leaves. The reference proteins of *N. benthamiana* were download from Sol Genomics Network database (https://solgenomics.net/ftp/genomes/Nicotiana_benthamianaV261/Nbenthamiana_Annotation/)

**Table S2.** Information on RNA-binding polypeptide (RBP) and RNA aptamers. Red font indicates shared amino-acid residues between ARM and RRRG degron.

| Name | Sequences with degon (RRRG) | RBP | Sequence | RNA ligand | Reference |
| --- | --- | --- | --- | --- | --- |
| Rev-RRRG | TRQARRNRRRRWRE<u>RRRG</u> | HIV-1 Rev | TRQARRNRRRRWRERQR | RRE | (8) |
| P22N-RRRG | NAKTRRHERRRKLAIE<u>RRRG</u> | P22N | NAKTRRHERRRKLAIER | P22 boxB | (9) |
| λN-RRRG | MDAQTRRRE<u>RRRG</u> | λN | MDAQTRRRERRAEKQAQWKAAN | boxB | (10) |
| BTat-RRRG | SGRSRRSRGTRGKGRRIR<u>RRG</u> | BIV-Tat | SGRSRRSRGTRGKGRRIRR | Pepper | (11) |
| HTat-RRRG | SYGRKKRRQ<u>RRRG</u> | HIV-Tat | SYGRKKRRQRRRRSRSQ | HIV-1 TAR | (12) |
| Rex-RRRG | MPKTRRRPRRSQRK<u>RRRG</u> | HTLV-1 Rex | MPKTRRRPRRSQRKRP | XBE | (13) |

**Table S3.**
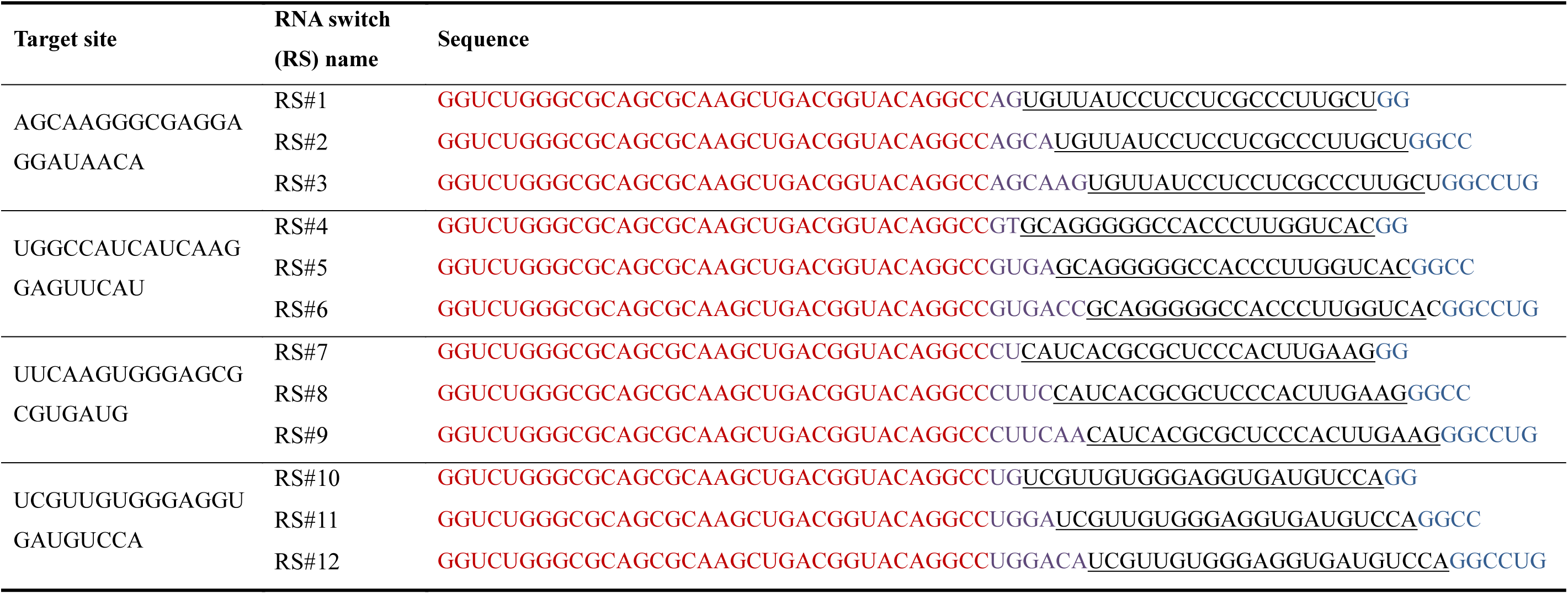
RNA switches for imaging *mCherry* RNA. The red, brown, black and blue font indicates respectively the RRE aptamer, linker 1, the probe sequence, and linker 2 in RNA switches.

**Table S4.**
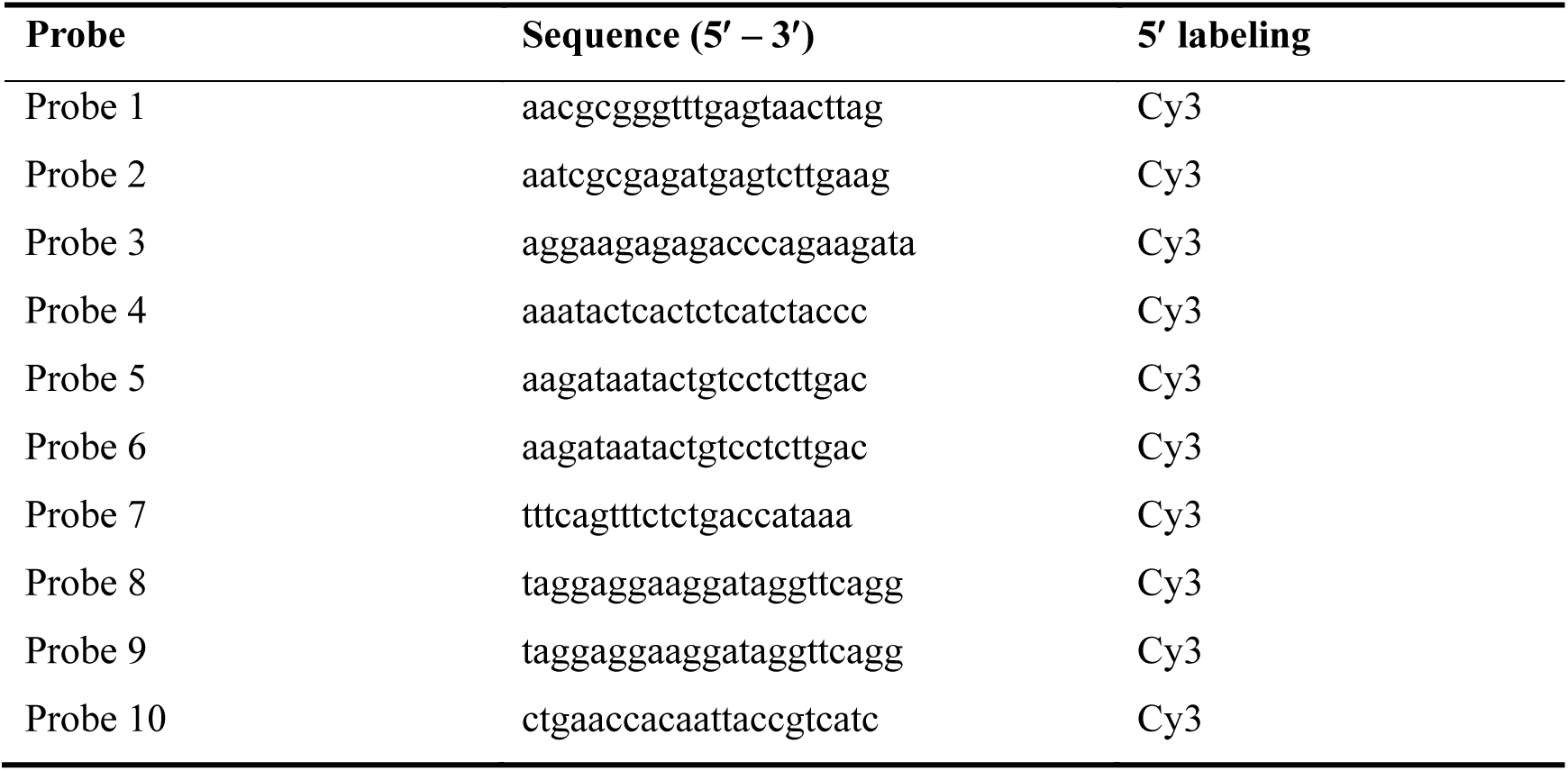
smFISH probes used for targeting *CM3* mRNA.

**Table S5.**
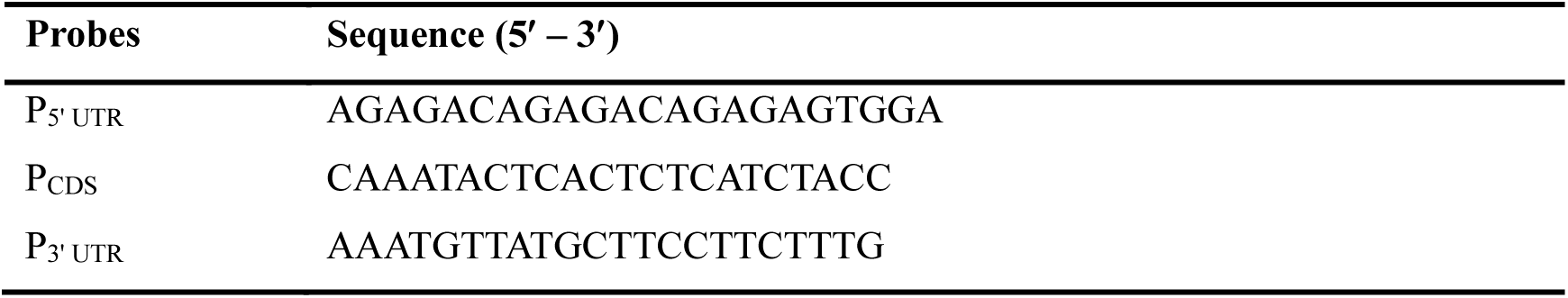
Specific probe sequences targeting the 5’ UTR, coding sequence and 3’ UTR of *CM3* mRNA.

**Table S6.** Identifiers for Arabidopsis target genes.

| Gene name | Locus | Gene ID |
| --- | --- | --- |
| *TUB5* | AT1G20010 | 838590 |
| *CCA1* | AT2G46830 | 819296 |
| *CIPK11* | AT2G30360 | 817586 |
| *SMB* | AT1G79580 | 844296 |

**Table S7.**
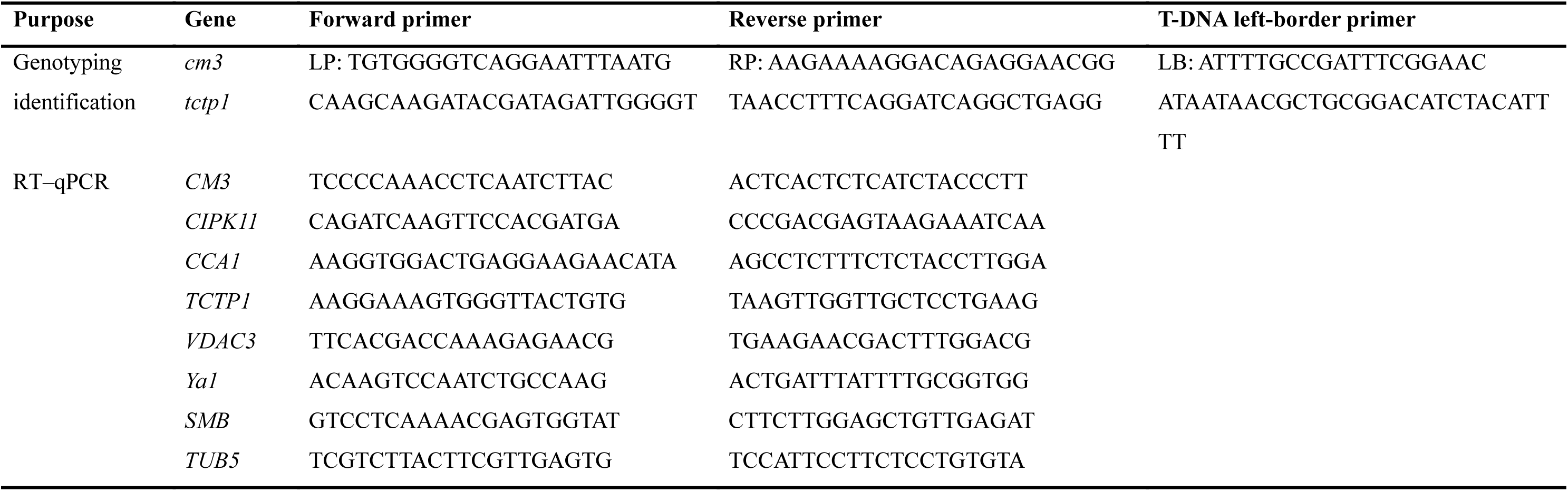
Primers for genotyping of Arabidopsis mutants and RT–qPCR.

**Table S8.**
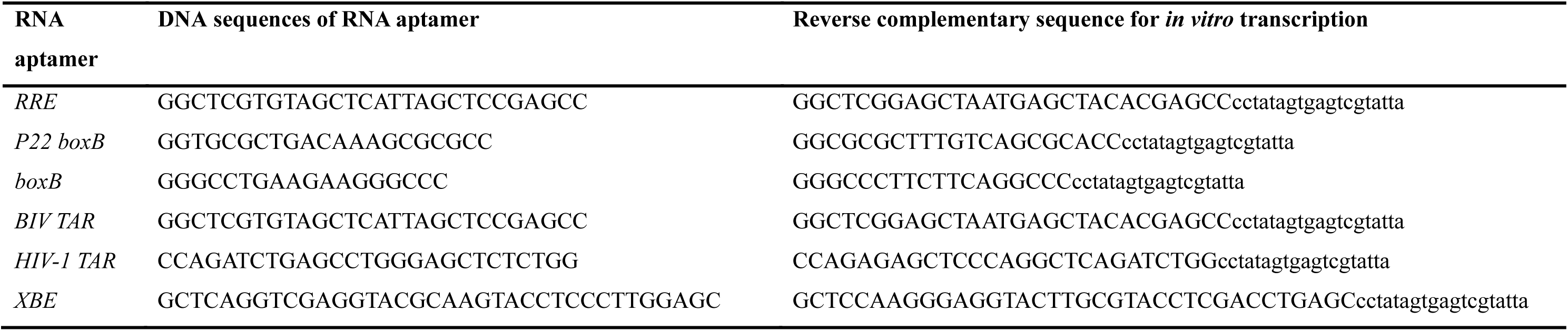
DNA sequences for *in vitro* transcription of RNA aptamers. Lowercase letters indicate the reverse complementary sequence of the *T7* promoter.

**Table S9.**
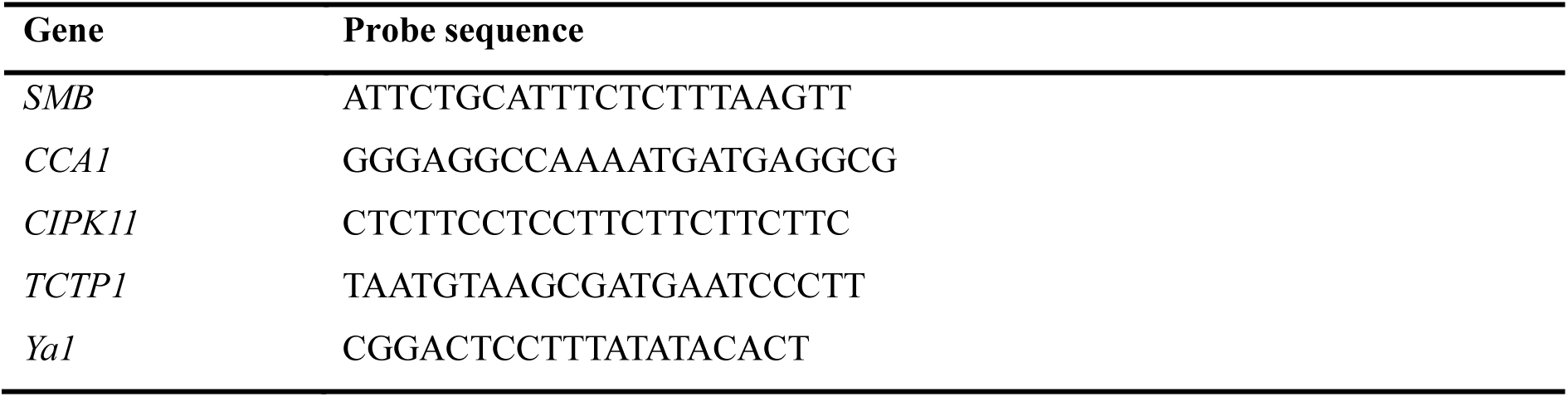
Probe sequences in RNA switch for imaging target gene expression.

**Table S10.**
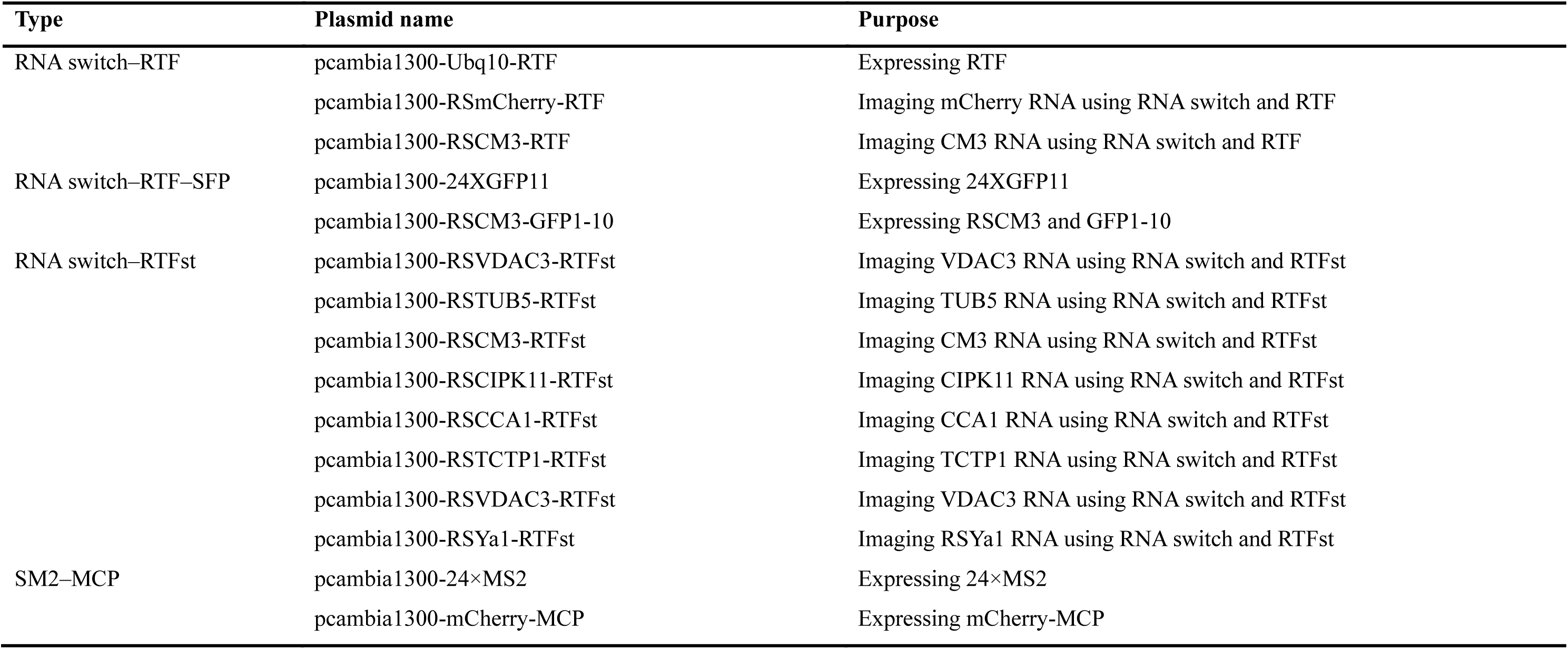
Plasmids used in this study.

| Type | Plasmid name | Purpose |
| --- | --- | --- |
| RNA switch–RTF | pcambia1300-Ubq10-RTF | Expressing RTF |
|  | pcambia1300-RSmCherry-RTF | Imaging mCherry RNA using RNA switch and RTF |
|  | pcambia1300-RSCM3-RTF | Imaging CM3 RNA using RNA switch and RTF |
| RNA switch–RTF–SFP | pcambia1300-24XGFP11 | Expressing 24XGFP11 |
|  | pcambia1300-RSCM3-GFP1-10 | Expressing RSCM3 and GFP1-10 |
| RNA switch–RTFst | pcambia1300-RSVDAC3-RTFst | Imaging VDAC3 RNA using RNA switch and RTFst |
|  | pcambia1300-RSTUB5-RTFst | Imaging TUB5 RNA using RNA switch and RTFst |
|  | pcambia1300-RSCM3-RTFst | Imaging CM3 RNA using RNA switch and RTFst |
|  | pcambia1300-RSCIPK11-RTFst | Imaging CIPK11 RNA using RNA switch and RTFst |
|  | pcambia1300-RSCCA1-RTFst | Imaging CCA1 RNA using RNA switch and RTFst |
|  | pcambia1300-RSTCTP1-RTFst | Imaging TCTP1 RNA using RNA switch and RTFst |
|  | pcambia1300-RSVDAC3-RTFst | Imaging VDAC3 RNA using RNA switch and RTFst |
|  | pcambia1300-RSYa1-RTFst | Imaging RSYa1 RNA using RNA switch and RTFst |
| SM2–MCP | pcambia1300-24×MS2 | Expressing 24×MS2 |
|  | pcambia1300-mCherry-MCP | Expressing mCherry-MCP |

Video S1.

Cell-to-cell trafficking of *TCTP1* mRNA in Arabidopsis root.

Video S2.

Visualization of aphid-secreted *Ya1* in *N. benthamiana* leaves.

## References

1. Hani, S. et al. Live single-cell transcriptional dynamics via RNA labelling during the phosphate response in plants. Nat. Plants 7, 1050–1064 (2021).

2. López-Maury, L., Marguerat, S., and Bähler, J. Tuning gene expression to changing environments: from rapid responses to evolutionary adaptation. Nat. Rev. Genet. 9, 583–593 (2008).

3. Huang, N. C., Luo, K. R., and Yu, T. S. Development of a split fluorescent protein-based RNA live-cell imaging system to visualize mRNA distribution in plants. Plant Methods 18, 1–10 (2022).

4. Alamos, S. et al. Quantitative imaging of RNA polymerase II activity in plants reveals the single-cell basis of tissue-wide transcriptional dynamics. Nat. Plants 7, 1037–1049 (2021).

5. Kitagawa, M., Tran, T. M., and Jackson, D. Traveling with purpose: cell-to-cell transport of plant mRNAs. Trends Cell Biol. 34, 48–57 (2024).

6. Liu, L. and Chen, X. Intercellular and systemic trafficking of RNAs in plants. Nat. Plants 4, 869–878 (2018).

7. Zhang, W. et al. Graft-transmissible movement of inverted-repeat-induced siRNA signals into flowers. Plant J. 80, 106–121 (2014).

8. Molnar, A. et al. Small silencing RNAs in plants are mobile and direct epigenetic modification in recipient cells. Science 328, 872–875 (2010).

9. Corbesier, L. et al. FT protein movement contributes to long-distance signaling in floral induction of Arabidopsis. Science 316, 1030–1033 (2007).

10. Han, W. H. et al. A small RNA effector conserved in herbivore insects suppresses host plant defense by cross-kingdom gene silencing. Mol. Plant 18, 437–456 (2025).

11. Zhang, Z. L., et al. Cross-kingdom RNA interference mediated by insect salivary microRNAs may suppress plant immunity. Proc. Natl. Acad. Sci. USA 121, e2318783121 (2024).

12. Westwood, J. H. and Kim, G. RNA mobility in parasitic plant - host interactions. RNA Biol. 14, 450–455 (2017).

13. van Kleeff, P. J. et al. Small RNAs from Bemisia tabaci Are Transferred to Solanum lycopersicum Phloem during Feeding. Front Plant Sci. 7, 1–12 (2016).

14. Wang, M., Thomas, N., and Jin, H. Cross-kingdom RNA trafficking and environmental RNAi for powerful innovative pre- and post-harvest plant protection. Curr. Opin. Plant Biol. 38, 133–141 (2017).

15. Bai, J. et al. A protein-independent fluorescent RNA aptamer reporter system for plant genetic engineering. Nat. Commun. 11, 3847 (2020).

16. Chen, X. J. et al. Visualizing RNA dynamics in live cells with bright and stable fluorescent RNAs. Nat. Biotechnol. 37, 1287–1293 (2019).

17. Haimovich, G. et al. Use of the MS2 aptamer and coat protein for RNA localization in yeast: A response to “MS2 coat proteins bound to yeast mRNAs block 5′ to 3′ degradation and trap mRNA decay products: implications for the localization of mRNAs by MS2-MCP system”. RNA 22, 660–666 (2016).

18. Garcia, J. F. and Parker, R. Ubiquitous accumulation of 3′ mRNA decay fragments in mRNAs with chromosomally integrated MS2 arrays. RNA 22, 657–659 (2016).

19. Garcia, J. F. and Parker, R. MS2 coat proteins bound to yeast mRNAs block 5′ to 3′ degradation and trap mRNA decay products: implications for the localization of mRNAs by MS2-MCP system. RNA 21, 1393–1395 (2015).

20. Park, H. Y. et al. Visualization of Dynamics of Single Endogenous mRNA Labeled in Live Mouse. Science 343, 422–424 (2014).

21. Göhring, J., Jacak, J., and Barta, A. Imaging of Endogenous Messenger RNA Splice Variants in Living Cells Reveals Nuclear Retention of Transcripts Inaccessible to Nonsense-Mediated Decay in Arabidopsis. Plant Cell 26, 754–764 (2014).

22. Wang, Z. J. et al. In Situ Spatial Complementation of Aptamer-Mediated Recognition Enables Live-Cell Imaging of Native RNA Transcripts in Real Time. Angewandte Chemie-International Edition 57, 972–976 (2018).

23. Xia, C. et al. Single-molecule live-cell RNA imaging with CRISPR-Csm. Nat Biotechnol. 1, 1–21 (2025).

24. Colognori, D., Trinidad, M., and Doudna, J. A. Precise transcript targeting by CRISPR-Csm complexes. Nat Biotechnol. 41, 1256–1264 (2023).

25. Yang, L. Z. et al. Dynamic Imaging of RNA in Living Cells by CRISPR-Cas13 Systems. Mol. Cell 76, 981–997 (2019).

26. Nelles, D. A. et al. Programmable RNA Tracking in Live Cells with CRISPR/Cas9. Cell 165, 488–496 (2016).

27. Le, P., Ahmed, N., and Yeo, G. W. Illuminating RNA biology through imaging. Nat. Cell Biol. 24, 815–824 (2022).

28. Liu, Y. et al. A split ribozyme system for in vivo plant RNA imaging and genetic engineering. Plant Biotechnol. J. 1, 1–10 (2025).

29. Yang, L. et al. m^5^C Methylation Guides Systemic Transport of Messenger RNA over Graft Junctions in Plants. Curr. Biol. 29, 2465–2476 (2019).

30. Chen, Y. Z. et al. An aphid RNA transcript migrates systemically within and is a virulence factor. P. Natl. Acad. Sci. USA 117, 12763–12771 (2020).

31. Chen, X. Y. et al. Molecular basis for arginine C-terminal degron recognition by Cul2^FEM1^ E3 ligase. Nat. Chem. Biol. 17, 1552–4450 (2021).

32. Sepers, J. J. et al. The mIAA7 degron improves auxin-mediated degradation in Caenorhabditiselegans. G3 (Bethesda) 12, 1-15 (2022).

33. Yesbolatova, A. et al. Generation of conditional auxin-inducible degron (AID) cells and tight control of degron-fused proteins using the degradation inhibitor auxinole. Methods 164, 73–80 (2019).

34. Austin, R. J. et al. Designed arginine-rich RNA-binding peptides with picomolar affinity. J. Am. Chem. Soc., 10966–10967 (2002).

35. Smith, C. A., Calabro, V., and Frankel, A. D. An RNA-Binding Chameleon. Mol. Cell 6, 1067–1076 (2000).

36. Jiang, F. et al. Anchoring an extended HTLV-1 Rex peptide within an RNA major groove containing junctional base triples. Structure, 1461–S12 (1999).

37. Legault, P. et al. NMR Structure of the Bacteriophage λ N Peptide/boxB RNA Complex: Recognition of a GNRA Fold by an Arginine-Rich Motif. Cell 93, 289–299 (1998).

38. Battiste, J. L. et al. Alpha helix-RNA major groove recognition in an HIV-1 rev peptide-RRE RNA complex. Science 273, 1547–1551 (1996).

39. Puglisi, J. D. et al. Solution Structure of a Bovine Immunodeficiency Virus Tat-TAR Peptide-RNA Complex. Science 270, 1200–1203 (1995).

40. Chen, M. et al. Live imaging of RNA and RNA splicing in mammalian cells via the dcas13a-SunTag-BiFC system. Biosens. Bioelectron. 204, 1–11 (2022).

41. Luo, K., Huang, N., and Yu, T. Selective Targeting of Mobile mRNAs to Plasmodesmata for Cell-to-Cell Movement. Plant Physiol., 604–614 (2018).

42. Biayna, J. and Dumbović, G. Decoding subcellular RNA localization one molecule at a time. Genome Biol 26, 1–23 (2025).

43. Fischer, A. et al. Membrane localization of acetylated CNK1 mediates a positive feedback on RAF/ERK signaling. Sci. Adv. 3, 1–10 (2017).

44. Michaud, M. et al. Differential targeting of VDAC3 mRNA isoforms influences mitochondria morphology. P. Natl. Acad. Sci. USA 111, 8991–8996 (2014).

45. Bach-Pages, M. et al. Discovering the RNA-Binding Proteome of Plant Leaves with an Improved RNA Interactome Capture Method. Biomolecules 10, 1–21 (2020).

46. Zheng, L. L. et al. A root cap-localized NAC transcription factor controls root halotropic response to salt stress in Arabidopsis. Nat. Commun. 15, 1–12 (2024).

47. de los Reyes, P. et al. CONSTANS alters the circadian clock in Arabidopsis thaliana. Mol. Plant 17, 1204–1220 (2024).

48. Gu, S. B. et al. The kinase CIPK11 functions as a positive regulator in cadmium stress response in Arabidopsis. Gene 772, 1–9 (2021).

49. Kondhare, K. R., Patil, N. S., and Banerjee, A. K. A historical overview of long-distance signalling in plants. J. Exp. Bot. 72, 4218–4236 (2021).

50. Wang, S. et al. Plant mRNAs move into a fungal pathogen via extracellular vesicles to reduce infection. Mol. Plant Microbe In. 37, 29–29 (2024).

51. Braselmann, E. et al. Illuminating RNA Biology: Tools for Imaging RNA in Live Mammalian Cells. Cell Chem. Biol. 27, 891–903 (2020).

52. Prigodich, A. E. et al. Nano-flares for mRNA Regulation and Detection. ACS Nano 3, 2147–2152 (2009).

53. Pham, T. G. and Wu, J. H. Recent advances in methods for live-cell RNA imaging. Nanoscale 16, 5537–5545 (2024).

54. Al Mazid, M. F. et al. Application of fluorescent turn-on aptamers in RNA studies. *Mol*. Omics 17, 483–491 (2021).

55. Truong, L. and Ferré-D’Amaré, A. R. From fluorescent proteins to fluorogenic RNAs: Tools for imaging cellular macromolecules. Protein Sci. 28, 1374–1386 (2019).

56. Singh, V. and Jain, M. Recent advancements in CRISPR-Cas toolbox for imaging applications. Crit. Rev. Biotechnol. 42, 508–531 (2022).

57. Chen, A. et al. Improving RNA-based crop protection through nanotechnology and insights from cross-kingdom RNA trafficking. Curr. Opin. Plant. Biol. 76, 1–22 (2023).

58. Weiberg, A. et al. Fungal Small RNAs Suppress Plant Immunity by Hijacking Host RNA Interference Pathways. Science 342, 118–123 (2013).

59. Fan, W. et al. Proteomics integrated with metabolomics: analysis of the internal causes of nutrient changes in alfalfa at different growth stages. BMC Plant Biol. 18, 1–15 (2018).

60. Demichev, V. et al. DIA-NN: neural networks and interference correction enable deep proteome coverage in high throughput. Nat. Methods 17, 41–44 (2020).

61. Yoo, S. D., Cho, Y. H., and Sheen, J. Arabidopsis mesophyll protoplasts: a versatile cell system for transient gene expression analysis. Nat. Protoc. 2, 1565–72 (2007).

## References

1. V. Demichev, C. B. Messner, S. I. Vernardis, K. S. Lilley, M. Ralser, DIA-NN: neural networks and interference correction enable deep proteome coverage in high throughput. Nat. Methods 17, 41–44 (2020).

2. S. J. Harrison et al., A rapid and robust method of identifying transformed seedlings following floral dip transformation. Plant Methods 2, 1–7 (2006).

3. N. Tsanov et al., smiFISH and FISH-quant - a flexible single RNA detection approach with super-resolution capability. Nucleic Acids Res. 44, 1–11 (2016).

4. M. Michaud et al., Differential targeting of VDAC3 mRNA isoforms influences mitochondria morphology. P. Natl. Acad. Sci. USA 111, 8991–8996 (2014).

5. C. Y. Li et al., Rice stripe virus activates the bZIP17/28 branch of the unfolded protein response signalling pathway to promote viral infection. Mol. Plant Pathol. 23, 447–458 (2022).

6. Z. Yang et al., RNase H1 Cooperates with DNA Gyrases to Restrict R-Loops and Maintain Genome Integrity in Arabidopsis Chloroplasts. Plant Cell 29, 2478–2497 (2017).

7. Y. S. Lai et al., Systemic signaling contributes to the unfolded protein response of the plant endoplasmic reticulum. Nat. Commun. 9, 1–11 (2018).

8. J. L. Battiste et al., Alpha helix-RNA major groove recognition in an HIV-1 Rev peptide-RRE RNA complex. Science 273, 1547–1551 (1996).

9. R. J. Austin, T. Xia, J. Ren, T. T. Takahashi, R. W. Roberts, Designed arginine-rich RNA-binding peptides with picomolar affinity. J. Am. Chem. Soc. 124, 10966–10967 (2002).

10. P. Legault, J. Li, J. Mogridge, L. E. Kay, J. Greenblatt, NMR Structure of the Bacteriophage λ N Peptide/boxB RNA Complex: Recognition of a GNRA Fold by an Arginine-Rich Motif. Cell 93, 289–299 (1998).

11. J. D. Puglisi, L. Chen, S. Blanchard, A. D. Frankel, Solution Structure of a Bovine Immunodeficiency Virus Tat-TAR Peptide-RNA Complex. Science 270, 1200–1203 (1995).

12. C. A. Smith, V. Calabro, A. D. Frankel, An RNA-Binding Chameleon. Mol. Cell 6, 1067–1076 (2000).

13. F. Jiang et al., Anchoring an extended HTLV-1 Rex peptide within an RNA major groove containing junctional base triples. Struct. Fold. Des. 7, 1461–1472 (1999).

